# Sensory cortex is optimised for prediction of future input

**DOI:** 10.1101/224758

**Authors:** Yosef Singer, Yayoi Teramoto, Ben D. B. WiĲmore, Andrew J. King, Jan W. H. Schnupp, Nicol S. Harper

**Affiliations:** Dept. of Physiology, Anatomy and Genetics (DPAG), Sherrington Building, University of Oxford, Parks Road, Oxford OX1 3PT, UK.; Dept. of Biomedical Science, City University of Hong Kong. 31 To Yuen Street, Kowloon Tong, Hong Kong.

## Abstract

Neurons in sensory cortex are tuned to diverse features in natural scenes. But what determines which features neurons become selective to? Here we explore the idea that neuronal selectivity is optimised to represent features in the recent past of sensory input that best predict immediate future inputs. We tested this hypothesis using simple feedforward neural networks, which were trained to predict the next few video or audio frames in clips of natural scenes. The networks developed receptive fields that closely matched those of real cortical neurons, including the oriented spatial tuning of primary visual cortex, the frequency selectivity of primary auditory cortex and, most notably, in their temporal tuning properties. Furthermore, the better a network predicted future inputs the more closely its receptive fields tended to resemble those in the brain. This suggests that sensory processing is optimised to extract those features with the most capacity to predict future input.

**Impact statement:** Prediction of future input explains diverse neural tuning properties in sensory cortex.

## Introduction

Sensory inputs guide actions, but such actions necessarily lag behind these inputs due to delays caused by sensory transduction, axonal conduction, synaptic transmission, and muscle activation. To strike a cricket ball, for example, one must estimate its future location, not where it is now. Prediction has other fundamental theoretical advantages: a system that parsimoniously predicts future inputs from their past, and that generalizes well to new inputs, is likely to contain representations that reflect their underlying causes^1^. This is important because ultimately we are interested in these causes (e.g. flying cricket balls), not the raw images or sound waves incident on the sensory receptors. Furthermore, much of sensory processing involves discarding irrelevant information, such as that which is not predictive of the future, to arrive at a representation of what is important in the environment for guiding action^1^.

Previous theoretical studies have suggested that many neural representations can be understood in terms of efficient coding of natural stimuli in a short time window at or just before the present^2–5^. Such studies generally built a network model of the brain, which was trained to represent stimuli subject to some set of constraints. One pioneering such study trained a network to efficiently represent static natural images using a sparse, generative model^4,5^. More recent studies have used related ideas to model the representation of moving (rather than static) images^6–8^ and other sensory stimuli^9–13^. In contrast, we built a network model that was optimised not for efficient representation of the recent past, but for efficient prediction of the immediate future of the stimulus, which we will refer to as the temporal prediction model.

To evaluate the representations produced by these theoretical models, we can compare them to the receptive fields of real neurons. Here, we define a neuron’s receptive field (RF) as the stimulus that maximally linearly drives the neuron. In mammalian primary visual cortex (V1), neurons typically respond strongly to oriented edge-like structures moving over a particular retinal location^14–17^ In mammalian primary auditory cortex (A1), most neurons respond strongly to changes in the amplitude of sounds within a certain frequency range^18^.

The temporal prediction model provides a principled approach to understanding the temporal aspects of RFs. Previous models, based on sparsity or slowness principles, were successful in accounting for many spatial aspects of V1 RF structure^4–8,19^, and had some success in accounting for spectral aspects of A1 RF structure^9–11,13^. However, these models do not account well for the temporal structure of V1 or A1 RFs. Notably, for both vision^17^ and audition^18^, the envelopes of real neuronal RFs tend to be asymmetric in time, with greater sensitivity to very recent inputs compared to inputs further in the past. In contrast, the RFs predicted by previous models typically show symmetrical temporal envelopes and lack a stronger sensitivity to the most recent past^6,9,10,12,13^.

Here we show that these shortcomings are largely overcome by the temporal prediction approach, suggesting that neural sensitivity at early levels of the cortical hierarchy may be organised to facilitate a rapid and efficient prediction of what the environment will look like in the next fraction of a second.

## Results

### The temporal prediction model

To determine what type of sensory RF structures would facilitate predictions of the imminent future, we built a feedforward network model with a single layer of nonlinear hidden units, mapping the inputs to the outputs through weighted connections (Fig. 1). Each hidden unit’s output results from a linear mapping (by input weights) from the past input, followed by a monotonic nonlinearity, much like the classic linear-nonlinear model of sensory neurons^9–11^. The model then generates a prediction of the future from a linear weighting of the hidden units’ outputs. This is consistent with the observation that decoding from the neural response is often well approximated by a linear transformation^20^.

**Figure 1.**
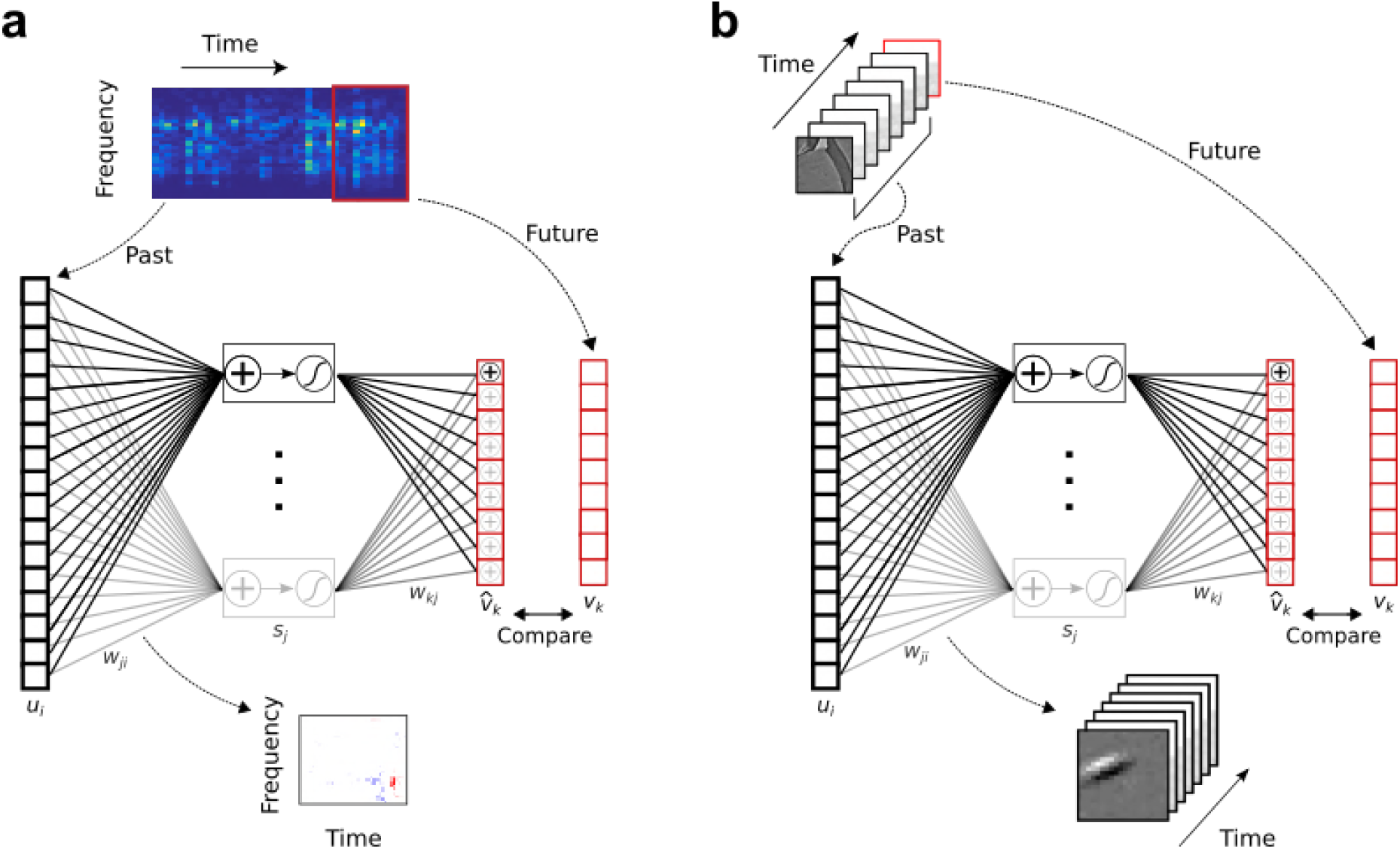
Temporal prediction model implemented using a feedforward artificial neural network, with the same architecture in both visual and auditory domains. **a,** Network trained on cochleagram clips (spectral content over time) of natural sounds, aims to predict immediate future time steps of each clip from recent past time steps. **b**, Network trained on movie clips of natural scenes, aims to predict immediate future frame of each clip from recent past frames. *u_i_*, input – the past; *w_ji_*, input weights; *s_j_*, hidden unit output; *w_kj_* output weights; 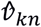, output – the predicted future; *v_k_*, target output – the true future. Hidden unit’s RF is the *w_ji_* between the input and that unit *j*.

We trained the temporal prediction model on extensive corpora, either of soundscapes or silent movies, modelling A1 (Fig. 1a) or V1 (Fig. 1b) neurons, respectively. In each case, the networks were trained by optimising their synaptic weights to most accurately predict the immediate future of the stimulus from its very recent past. For vision, the inputs were patches of videos of animals moving in natural settings, and we trained the network to predict the pixel values for one movie frame (40 ms) into the future, based on the 7 most recent frames (280 ms). For audition, we trained the network to predict the next three time steps (15 ms) of cochleagrams of natural sounds based on the 40 most recent time steps (200 ms). Cochleagrams resemble spectrograms but are adjusted to approximate the auditory nerve representation of sounds (see Methods).

During training we used sparse, *L*_1_ weight regularisation (see Eqn. 3 in Methods) to constrain the network to predict future stimuli in a parsimonious fashion, forcing the network to use as few weights as possible while maintaining an accurate prediction. This constraint can be viewed as an assumption about the sparse nature of causal dependencies underlying the sensory input, or alternatively as analogous to the energy and space restrictions of neural connectivity. It also prevents our network model from overfitting to its inputs.

### Qualitative assessment of auditory receptive fields

To compare with the model, we recorded responses of 114 auditory neurons (including 76 single units) in A1 and the anterior auditory field of 5 anesthetised ferrets^21^ and measured their spectrotemporal RFs (see Methods). The RFs are diverse (Fig. 2a); their frequency tuning can be narrowband, broadband or more complex, sometimes showing flanking inhibition. They may also lack clear order or be selective for the direction of frequency modulation^22^.

**Figure 2.**
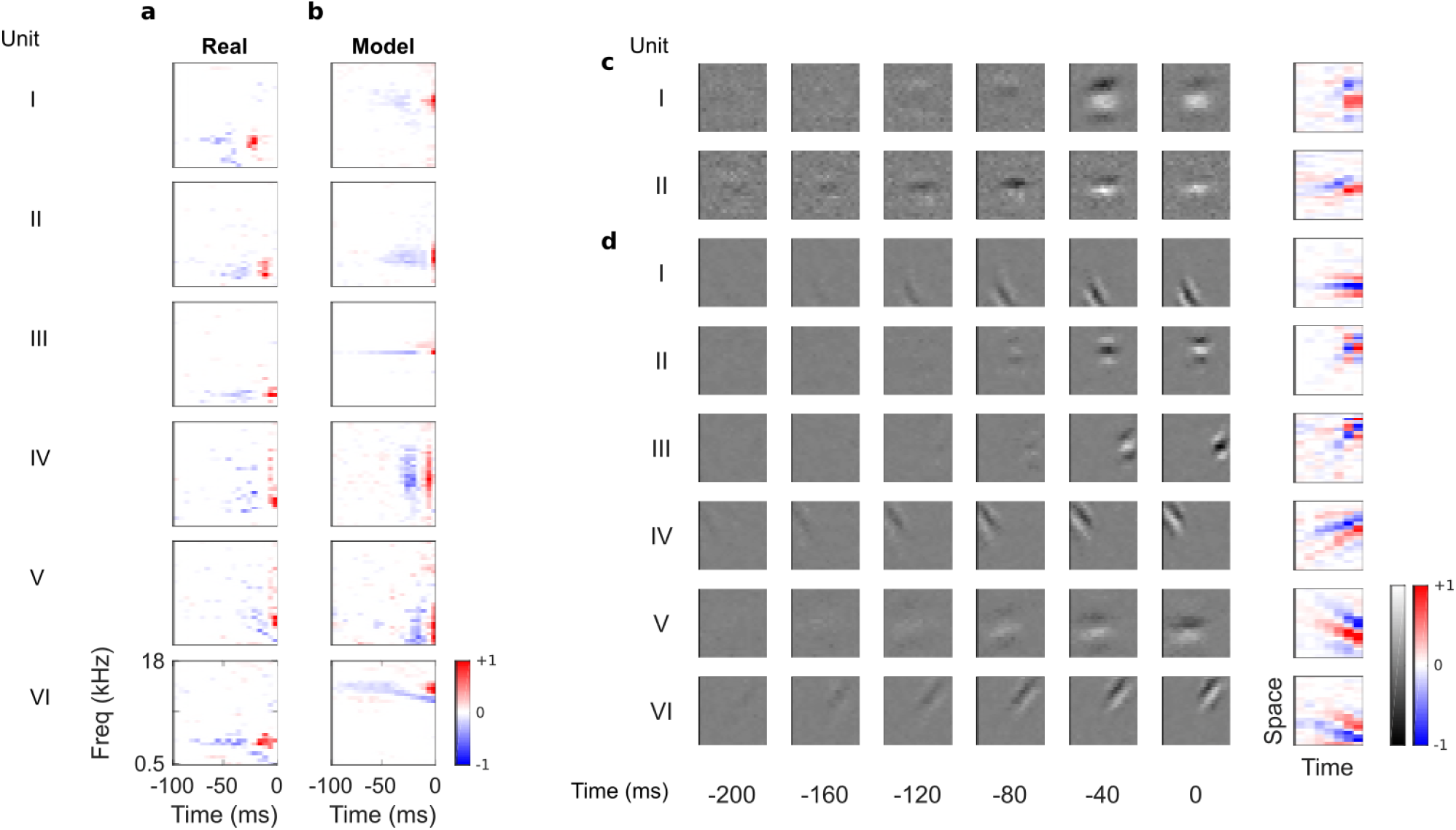
Auditory spectrotemporal and visual spatiotemporal RFs of real neurons and temporal prediction model units. **a,** Example spectrotemporal RFs of real A1 neurons^21^. Red – excitation, blue – inhibition. **b,** Example spectrotemporal RFs of model units when model is trained to predict the future of natural sound inputs. **c,** Example spatiotemporal RFs of real V1 neurons. Left, grayscale: 3D (space-space-time) spatiotemporal RFs showing the spatial RF at each of the most recent 6 timesteps. White – excitation, black – inhibition. Right: corresponding 2D (space-time) spatiotemporal RFs obtained by summing along the unit’s axis of orientation for each timestep. Red – excitation, blue – inhibition. **d,** Example 3D and corresponding 2D spatiotemporal RFs of model units when model is trained to predict the future of natural visual inputs.

In their temporal tuning, A1 RFs tend to weight recent inputs more heavily, with a temporally asymmetric power profile, involving excitation near the present followed by lagging inhibition of a longer duration^18^. The temporal prediction model RFs (Fig. 2b) are similarly diverse, showing all of the RF types seen *in vivo* (including examples of localised, narrowband, broadband, complex, disordered and directional RFs) and are well matched in scale and form to those measured in A1. This includes having greater power (root mean square) near the present, with brief excitation followed by longer lagging inhibition, producing an asymmetric power profile. This stands in contrast to previous attempts to model RFs based on efficient and sparse coding hypotheses, which either did not capture the diversity of RFs^11^, or lacked temporal asymmetry, punctate structure, or appropriate time scale^9,10,12,13,22,23^.

### Qualitative assessment of visual receptive fields

We also found substantial similarities when we compared the temporal prediction model’s RFs trained using visual inputs (Fig. 1b) with the 3D (space-space-time) and 2D (space-time) spatiotemporal RFs of real V1 simple cells, which were obtained from Ohzawa et al^24^. Simple cells^14^ have stereotyped RFs containing parallel, spatially localised excitatory and inhibitory regions, with each cell having a particular preferred orientation and spatial frequency^15–17^ (Fig. 2c). These features are also clearly apparent in the model RFs (Fig. 2d).

Unlike previous models^6,25,26^, the temporal prediction model captures the temporal asymmetry of real RFs. The RF power is highest near the present and decays exponentially into the past (Fig. 2d), as observed in real neurons^24^ (Fig. 2c). Furthermore, simple cell RFs have two types of spatiotemporal structure: space-time separable RFs (Fig. 2cI), whose optimal stimulus resembles a flashing or slowly ramping grating, and space-time inseparable RFs, whose optimal stimulus is a drifting grating^16^ (Fig. 2cII). Our model captures this diversity (Fig. 2dI-III separable, Fig. 2dIV-VI inseparable).

### Qualitative comparison to other models

For comparison, we trained a sparse coding model^4,5,10^(https://github.com/zayd/sparsenet) using our dataset. We would expect such a model to perform less well in the temporal domain, because unlike the temporal prediction model, the direction of time is not explicitly accounted for. The sparse coding model was chosen because it has set the standard for normative models of visual RFs^4–6,19^, and the same model has also been applied for auditory RFs^10,23,27,28^. Past studies^4,5,10^ have largely analysed the basis functions produced by the sparse coding model and compared their properties to neuronal RFs. To be consistent with these studies we have done the same, and to have a common term, refer to the basis functions as RFs (although strictly, they are projective fields). We can visually compare the large set of RFs recorded from A1 neurons (Fig. 3) to the full set of RFs obtained from the temporal prediction model when trained on auditory inputs (Fig. 4) and those of the sparse coding model (Fig. 5) when trained on the same auditory inputs.

**Figure 3.**
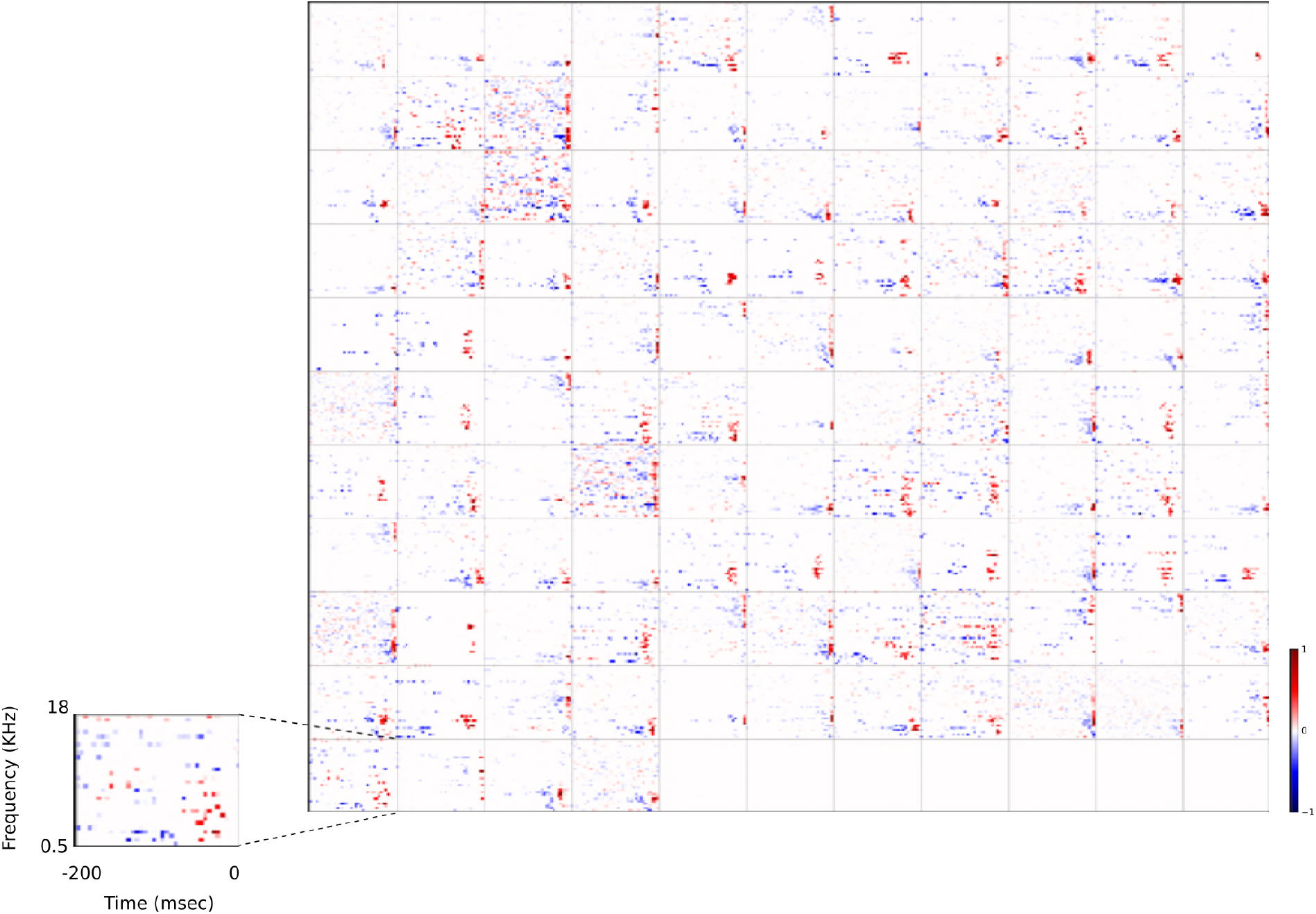
Full dataset of real auditory RFs. 114 neuronal RFs in A1 and AAF of 5 ferrets. Red – excitation, blue – inhibition. Inset shows axes.

**Figure 4.**
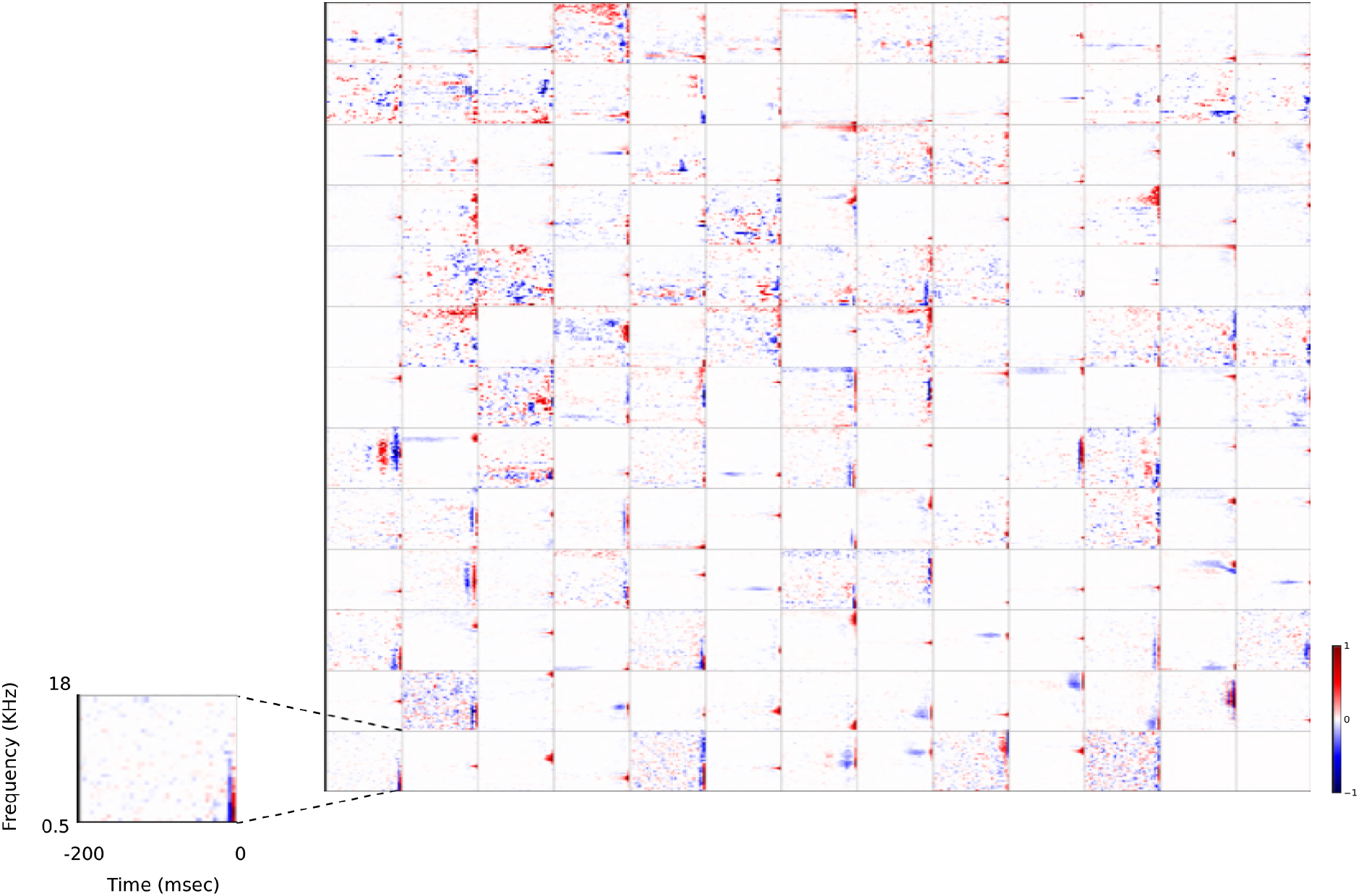
Full set of auditory RFs of the temporal prediction model units. Units were obtained by training the model with 1600 hidden units on auditory inputs. The hidden unit number and L1 weight regularization strength (10^−3.5^) was chosen because it results in the lowest MSE on the prediction task, as measured using a cross validation set. Many hidden units’ weight matrices decayed to near zero during training (due to the L1 regularization), leaving 167 active units. Inactive units were excluded from analysis and are not shown. Example units in Figure 2 come from this set. Red – excitation, blue – inhibition. Inset shows axes. Figure Supplement 1 shows the same RFs on a finer timescale. The full sets of visual spatial and corresponding spatiotemporal RFs for the temporal prediction model when it is trained on visual inputs are shown in Figure Supplements 2-3. Figure Supplement 4 shows the auditory RFs of the temporal prediction model when a linear activation function instead of a sigmoid nonlinearity was used. Figure Supplements 5-7 show the auditory spectrotemporal and visual spatial and 2D spatiotemporal RFs of the temporal prediction model when it was trained on inputs without added noise.

**Figure 5.**
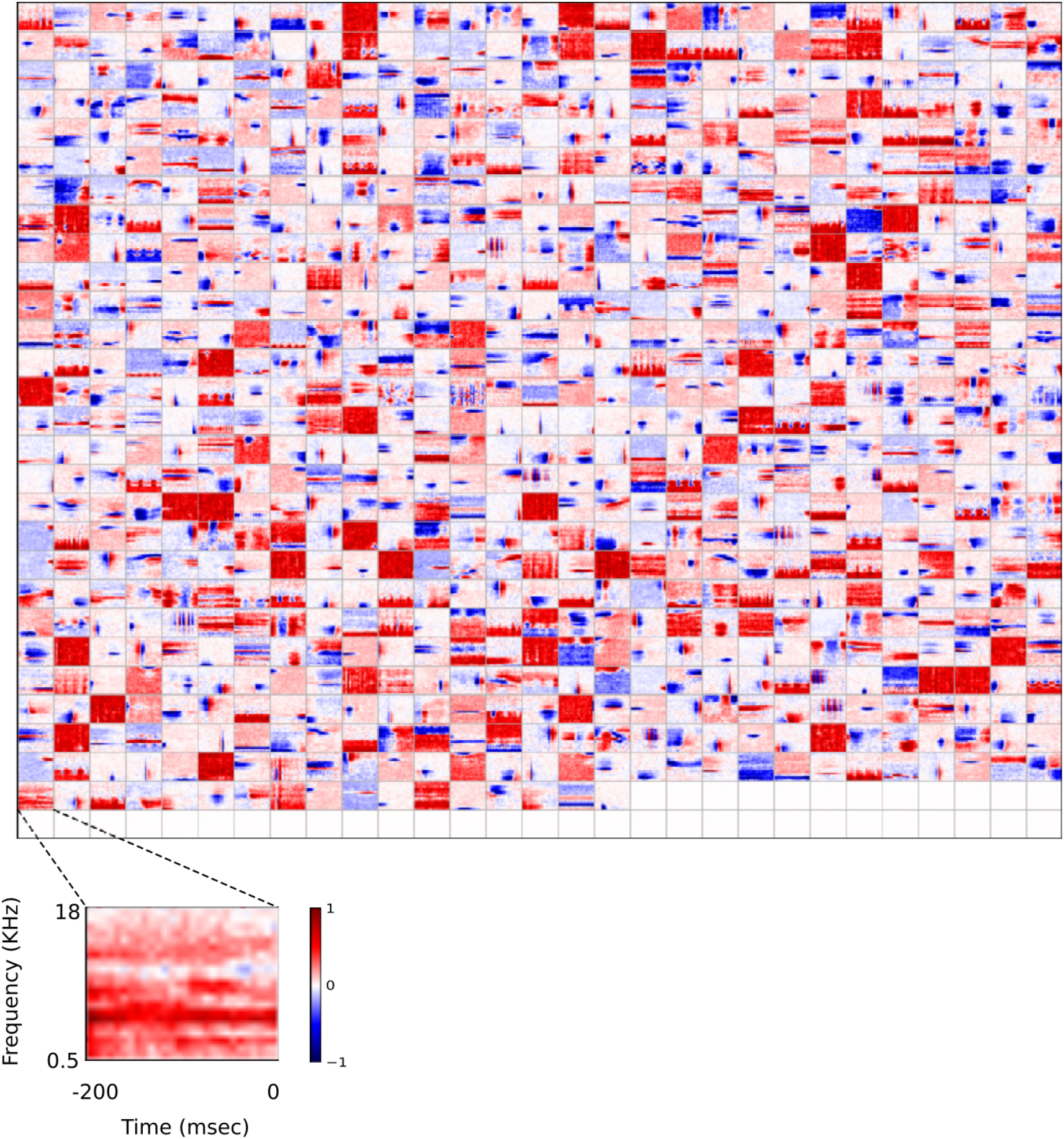
Full set of auditory ‘RFs’ (basis functions) of sparse coding model used as a control. Units were obtained by training the sparse coding model with 800 units on the identical auditory inputs used to train the network shown in Figure 4. L1 regularization of 10^0.5^ was applied to the hidden units’ activities. This network configuration was selected as it produced unit RFs that most closely resembled those recorded in A1, as determined by visual inspection. Although the basis functions of the sparse coding model are not receptive fields, but projective fields, they tend to be similar in structure^4,5^. In this manuscript, to have a common term between models and the data, we refer to sparse coding basis functions as RFs. Red – excitation, blue – inhibition. Inset shows axes. The full sets of visual spatial and corresponding spatiotemporal RFs for the sparse coding model when it is trained on visual inputs are shown in Figure Supplements 1-2. Figure Supplements 3-5 show the auditory spectrotemporal and visual spatial and 2D spatiotemporal RFs of the sparse coding model when it was trained on inputs without added noise

A range of RFs were produced by the sparse coding model, some of which show characteristics reminiscent of A1 RFs, particularly in the frequency domain. However, the temporal properties of A1 neurons are not well captured by these RFs. While some RFs display excitation followed by lagging inhibition, very few, if any, show distinct brief excitation followed by extended inhibition. Instead, neurons that show both excitation and inhibition tend to have a symmetric envelope and these features are randomly localised in time, and many neurons display temporally elongated structures that are not found in A1 neurons.

As with the auditory model, we also trained the sparse coding model on the identical visual inputs to serve as a control. We compared the full population of spatial and 2D spatiotemporal visual RFs of the temporal prediction model (Fig. 4-Fig. Supplements 2–3) and the sparse coding model (Fig. 5-Fig. Supplements 1-2). As shown in previous studies^4–6,19^, the sparse coding model produces RFs whose spatial structure resembles that of V1 simple cells (Fig. 5-Fig. Supplements 1-2), but does not capture the asymmetric nature of the temporal tuning of V1 neurons. Furthermore, while it does produce examples of both separable and inseparable spatiotemporal RFs, those that are separable tend to be completely stationary over time, resembling immobile rather than flashing gratings (Fig. 5-Fig. Supplement 2).

We also examined variants of the temporal prediction model. When a different hidden unit nonlinearity (tanh) was used, the networks had similar predictive capacity and produced comparable RFs. However, when the temporal prediction model had linear hidden units, it no longer predicted as well and produced RFs that were less like real neurons in their structure, generally being narrowband in frequency with temporally extended excitation (Fig. 4 Fig. Supplement 4).

### Quantitative analysis of auditory results

We compared parameters of the RFs generated by both models to the RFs of the population of real A1 neurons we recorded. We first compared the RFs in a non-parametric manner, using multidimensional scaling to embed each RF in a two-dimensional plane (Fig. 6a). RFs from the real population and those from the temporal prediction model cluster tightly together in the centre of this plane. In contrast, there is little overlap between the locations of the sparse coding RFs in this space and those of the real neurons.

We then examined specific attributes of the RFs to determine points of similarity and difference between each of the models and the recorded data. We first considered the temporal properties of the RFs and found that for the data and the temporal prediction model, most of the power is contained in the most recent time-steps (Figs. 2a-b, 3–4, 6b, and Fig. 4-Fig. Supplement 1). Given that the direction of time is not explicitly accounted for in the sparse coding model, as expected, it does not show this feature (Figs. 5, 6b). Next, we examined the tuning widths of the RFs in each population for both time and frequency, looking at excitation and inhibition separately. In the time domain, the real data tend to show leading excitation followed by lagging inhibition of longer duration (Figs. 2a, 3, 6c-e). The temporal prediction model also shows many neurons with this temporal structure, with lagging inhibition of longer duration than the leading excitation (Figs. 2b, 4, 6c-e, and Fig. 4-Fig. Supplement 1). This is not the case with the sparse coding model, where units tend to show either excitation and inhibition having the same duration or an elongated temporal structure that does not show such stereotyped polarity changes (Figs. 5, 6c-e). It is also the case that the absolute timescales of excitation and inhibition match more closely in the case of the temporal prediction model (Fig. 6c-e), although a few units display inhibition of a longer duration than is seen in the data (Fig. 6c).

**Figure 6.**
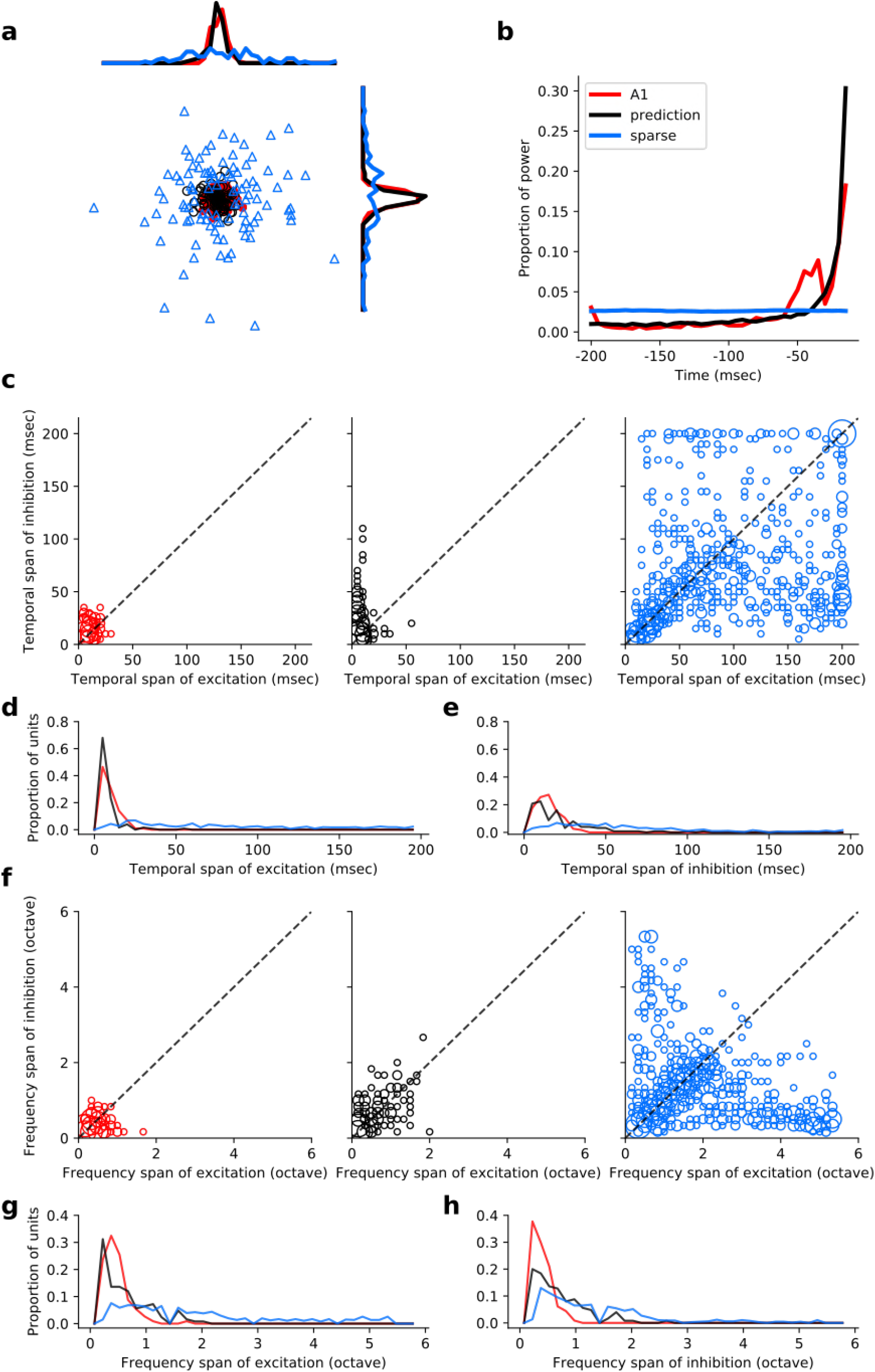
Population measures for real A1 plus temporal prediction and sparse coding model auditory spectrotemporal RFs. The population measures are taken from the RFs shown in Figures 3–5. **a,** Each point represents a single RF (with 32 frequency and 38 time bins) which has been embedded in a 2 dimensional space using Multi-Dimensional Scaling (MDS). Red circles – real A1 neurons, black circles – temporal prediction model units, blue triangles – sparse coding model units. Colour scheme applies to all subsequent panels. **b,** Proportion of power contained in each time step of the RF, taken as an average across the population of units. **c,** Temporal span of excitatory subfields versus that of inhibitory subfields, for real neurons and temporal prediction and sparse coding model units. The area of each circle is proportional to the number of occurrences at that point. **d,** Distribution of temporal spans of excitatory subfields, taken by summing along the x-axis in **c. e,** Distribution of temporal spans of inhibitory subfields, taken by summing along the y-axis in **c. f,** Frequency span of excitatory subfields versus that of inhibitory subfields, for real neurons and temporal prediction and sparse coding model units. **g,** Distribution of frequency spans of excitatory subfields, taken by summing along the x-axis in **f. h,** Distribution of frequency spans of inhibitory subfields, taken by summing along the y-axis in **f.** Figure Supplement 1 shows the same analysis for the temporal prediction model and sparse coding model trained on auditory inputs without added noise.

Regarding the spectral properties of real neuronal RFs, the spans of inhibition and excitation over sound frequency tend to be similar (Fig. 6f-h). This is also seen in the temporal prediction model, albeit with slightly more variation (Fig. 6f-h). The sparse coding model shows more extensive variation in frequency spans than either the data or our model (Fig. 6f-h).

### Quantitative analysis of visual results

We also compared the spatiotemporal RFs derived from the temporal prediction and sparse coding models with restricted published datasets summarizing RF characteristics of V1 neurons^17^ and a small number of full spatiotemporal visual RFs acquired from Ohzawa et al^24^. We assessed the orientation and spatial frequency tuning properties of the models’ RFs by fitting Gabor functions to their RFs (see Methods).

We compared temporal properties of the RFs from the neural data and the temporal prediction model. In both cases, most power (sum of square of RFs) is in the most recent time steps (Fig. 7a). Previous normative models of spatiotemporal RFs^6,25,26^ (Fig. 7-Fig. Supplement 1c-d) do not show this property, being either invariant over time or localised, but with a symmetric profile that is not restricted to the recent past. We also measured the space-time separability of the RFs of the temporal prediction model (see Methods); substantial numbers of both space-time separable and inseparable units were apparent (1038 separable, 455 inseparable; Fig. 4-Fig. Supplement 3). Finally, we observed an inverse correlation (r^2^ = -0.61, p < 10^−9^, n = 1098) between temporal and spatial frequency tuning (See Methods), which is also a property of real V1 RFs^16^ and is seen in a sparse-coding-related model^6^.

**Figure 7.**
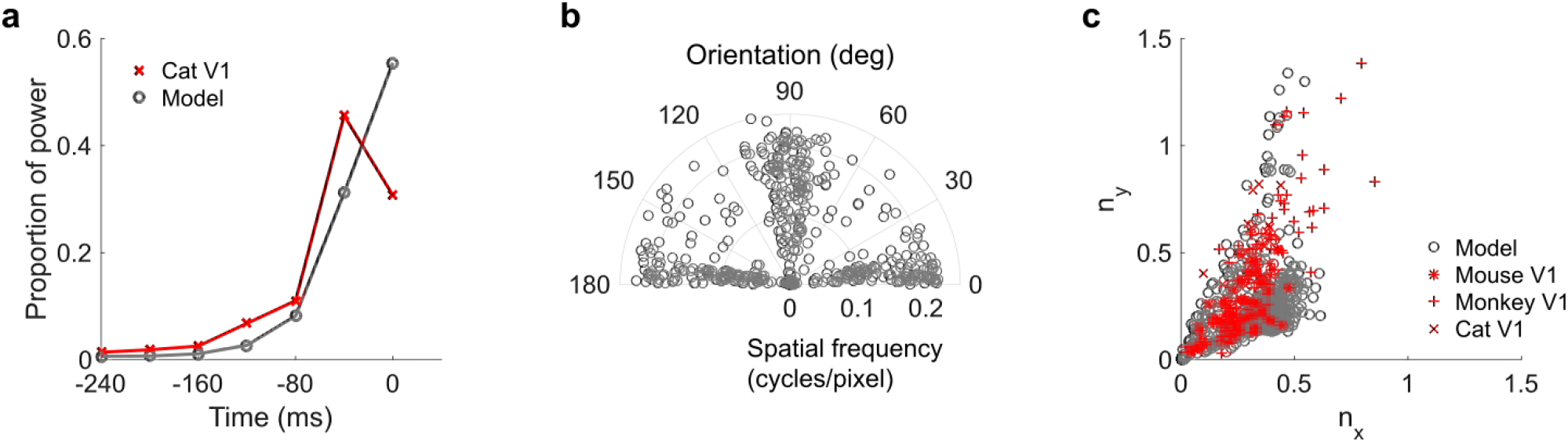
Population measures for real V1 and temporal prediction model visual spatial and spatiotemporal RFs. Model units were obtained by training the model with 1600 hidden units on visual inputs. The hidden unit number and L1 weight regularization strength (10^−3.75^) was chosen because it results in the lowest MSE on the prediction task, as measured using a cross validation set. Some hidden units’ weight matrices decayed to near zero during training (due to the L1 regularization), leaving 1493 active units. Inactive units were excluded from analysis. Example units in Figure 2 come from this set. **a,** Proportion of power (sum of squared weights over space and averaged across units) in each time step, for real^24^ and model populations. **b,** Joint distribution of spatial frequency and orientation tuning for population of model unit RFs at their timestep with greatest power. **c,** Distribution of RF shapes for real neurons (cat^15^, mouse^30^ and monkey^17^) and model units. *n_x_* and *n_y_* measure RF span parallel and orthogonal to orientation tuning, as a proportion of spatial oscillation period^17^. For **b-c**, only units that could be well approximated by Gabor functions (n = 1098 units; see Methods) were included in the analysis. Of these, only model units that were space-time separable (n = 677) are shown in **c** to be comparable with the neuronal data^17^. Figure Supplements 1-3 show example visual RFs and the same population measures for the sparse coding model trained on visual inputs with added noise and for the temporal prediction and sparse coding models trained on visual inputs without added noise.

The spatial tuning characteristics of the temporal prediction model’s RFs displayed a wide range of orientation and spatial frequency preferences, consistent with the neural data^16,29^ (Fig. 7b, Fig. 4-Fig. Supplement 2). Both model and real RFs^26^ show a preference for spatial orientations along the horizontal and vertical axes, although this orientation bias is seen to a greater extent in the temporal prediction model than in the data. The orientation and frequency tuning characteristics are also well captured by sparse coding related models of spatiotemporal RFs^6,25^ (Fig. 7-Fig. Supplement 1e). Furthermore, the widths and lengths of the RFs of the temporal prediction model, relative to the period of their oscillation, also match the neural data well (Fig. 7c). Although this property is again fairly well captured by previous model^4,5,8,17,19^ (Fig. 7-Fig. Supplement 1f), only the temporal prediction model seems to be able to capture the blob-like RFs that form a sizeable proportion of the neural data^17^ (Fig.7c where *n*_x_ and *n*_y_ < ~0.25, Fig. 4-Fig. Supplement 2).

### Optimising predictive capacity

Under our hypothesis of temporal prediction, we would expect that the better the temporal prediction model network is at predicting the future, the more the RFs of the network should resemble those of real neurons. To examine this hypothesis, we plotted the prediction error of the network as a function of two hyperparameters; the regularisation strength and the number of hidden units (Fig. 8a). Then, we plotted the similarity between the auditory RFs of real A1 neurons and those of the temporal prediction model (Fig. 8b), as measured by the mean KS distances of the temporal and frequency span distributions (Fig. 6d-e, g-h, Methods). The set of hyperparameter settings that give good predictions are also those where the temporal prediction model produces RFs that are most similar to those recorded in A1 (r^2^ = 0.8, p < 10^−9^, n = 55). This result argues that cortical neurons are indeed optimised for temporal prediction.

**Figure 8.**
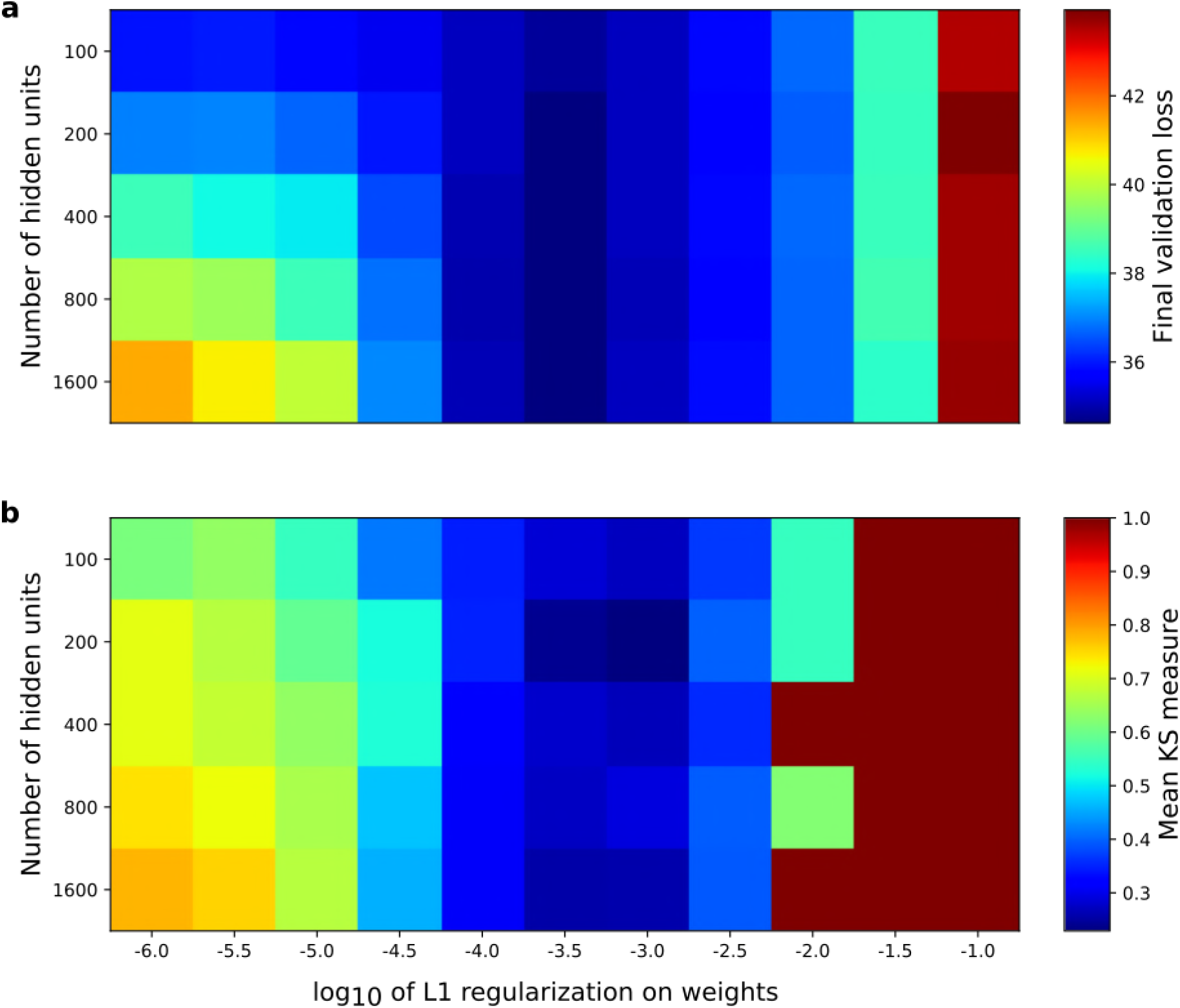
Correspondence between the temporal prediction model’s ability to predict future auditory input and the similarity of its units’ responses to those of real A1 neurons. Performance of model as a function of number of hidden units and regularization on the weights as measured by **a,** prediction error on validation set at the end of training and **b,** similarity between model units and real A1 neurons. The similarity between the real and model units is measured by averaging the Kolmogorov-Smirnov distance between each of the real and model distributions for the span of temporal and frequency tuning of the excitatory and inhibitory RF subfields (e.g. the distributions in Fig. 6d-e and Fig. 6g-h). Figure Supplement 1 shows the same analysis, performed for the sparse coding model, which does not produce a similar correspondence.

When the similarity measure was examined as a function of the same hyperparameters for the sparse coding model (Fig. 8-Fig. Supplement 1), and this was compared to that model’s stimulus reconstruction capacity as a function of the same hyperparameters, a monotonic relationship between stimulus reconstruction capacity and similarity of real RFs was not found (Fig. 8-Fig. Supplement 1; r^2^ = -0.05, p = 0.69, n = 50). In previous studies in which comparisons have been made between normative models and real data, the model hyperparameters have been selected to maximise the similarity between the real and model RFs. In contrast, the temporal prediction model provides an independent criterion, the prediction error, to perform hyperparameter selection.

## Discussion

We hypothesized that finding features that can efficiently predict future input from its past is a principle that influences the structure of sensory RFs. We implemented an artificial neural network model that instantiates a restricted version of this hypothesis. When this model was trained using natural sounds, it produced RFs that are both qualitatively and quantitatively similar to those of A1 neurons. Similarly, when we trained the model using natural movies it produced RFs with many of the properties of V1 simple cells. This similarity is particularly notable in the temporal domain; the model RFs have asymmetric envelopes, with a preference for the very recent past, as is seen in A1 and V1. Finally, the more accurate a temporal prediction model is at prediction, the more its RFs tend to be like real neuronal RFs.

### Relationship to other models

A number of principles, often acting together, have been proposed to explain the form and diversity of sensory RFs. These include efficient coding^3–5,10,11,23,25,31,32^, sparseness^4–6,10,12,19,23,25^, and slowness^22,26^. Efficient coding indicates that sensory input should be represented as accurately as possible given certain constraints, such as spike count or energy costs. Sparseness posits that only a small proportion of neurons in the population should be active for a given input. Finally, slowness means that neurons should be sensitive to features that change slowly over time. The temporal prediction principle we describe here provides another unsupervised objective of sensory coding. It has been described in a very general manner by the information bottleneck concept^1,33,34^. We have instantiated a specific version of this idea, with linear-nonlinear encoding of the input, followed by a linear transform from the encoding units’ output to the prediction.

In the following discussion, we describe previous normative models that infer RFs with temporal structure from auditory or movie input and relate them to spectrotemporal RFs in A1 or simple cell spatiotemporal RFs in V1, respectively. For focus, other normative models of less directly relevant areas, such as spatial receptive fields without a temporal component^4,5^, complex cells^7^, retinal receptive fields^32,35^, or auditory nerve impulse responses^36^, will not be examined.

### Auditory normative models

A number of coding objectives have been explored in normative models of A1 spectrotemporal RFs. One approach^11^ found analytically that the optimal typical spectrotemporal RF for efficient coding was spectrally localised with lagging and flanking inhibition, and showed an asymmetric temporal envelope. However, the resulting RF also showed substantially more flanking inhibition, more ringing over time and frequency, and operated over a much shorter timescale (~10ms) than seen in A1 RFs (Fig. 3). Moreover, this approach produced a single generic RF, rather than capturing the diversity of the population.

Other models have produced a diverse range of spectrotemporal RFs. In the sparse coding approach^10,23,27,28^, a spectrogram snippet is reconstructed from a sum of basis functions (a linear generative model), each weighted by its unit’s activity, with a constraint to have few active units. This approach is the same as the sparse coding model we used as a control (Fig. 5). A challenge with many sparse generative models is that the activity of the units is found by a recurrent iterative process that needs to find a steady state; this is fine for static stimuli such as images, but for dynamic stimuli like sounds it is questionable whether the nervous system would have sufficient time to settle on appropriate activities before the stimulus had changed. Related work also used a sparsity objective, but rather than minimising stimulus reconstruction error, forced high dispersal^12^ or decorrelation^9,22^ of neural responses. Although lacking some of the useful probabilistic interpretations of sparse generative models, this approach does not require a settling process for inference. An alternative to sparseness is temporal slowness, which can be measured by temporal coherence^22^. Here the linear transform from sequential spectrogram snippets to unit activity is optimised to maximise the correlation of each unit’s response over a certain time window, while maintaining decorrelation between the units’ activities.

Although the frequency tuning derived with these models can resemble that found in the midbrain or cortex^9,10,12,22,23,27,28^ (Fig. 5), the resulting RFs lack the distinct asymmetric temporal profile and lagging inhibition seen in real midbrain or A1 RFs. Furthermore, they often have envelopes that are too elongated over time, often spanning the full temporal width of the spectrotemporal RF. This is related to the fact that the time window to be encoded by the model is set arbitrarily, and every time point within that window is given equal weighting, i.e. the direction of time is not accounted for. This is in contrast to the temporal prediction model, which naturally gives greater weighting to time-points near the present than the past due to their greater predictive capacity.

### Visual normative models

The earliest normative model of spatiotemporal RFs of simple cells used independent component analysis (ICA)^6^, which is mathematically equivalent to the critically complete case of the sparse coding model^4,5^ we used as a control (Fig. 5-Fig. Supplements 1–2 and Fig. 7-Fig. Supplement 1). The RFs produced by this model and our control model reproduced fairly well the spatial aspects of simple cell RFs. However, in contrast to the temporal prediction model, the subset of more ‘blob-like’ RFs seen in the data are not well captured by our control sparse coding model (Fig. 7c and Fig. 7-Fig. Supplement 1f). In the temporal domain, again unlike the temporal prediction model and real V1 simple cells, the RFs of the ICA and sparse coding model are not pressed up against the present with an asymmetrical temporal envelope, but instead show a symmetrical envelope or span the entire range of times examined. A related model^25^ assumes that a longer sequence of frames is generated by convolving each basis function with a time-varying sparse coefficient and summing the result, so that each basis function is applied at each point in time. The resulting spatiotemporal RFs are similar to those produced by ICA^6^, or our control model (Fig. 5-Fig. Supplement 2 and Fig. 7-Fig. Supplement 1c). Although they tend not to span the entire range of times examined, they do show a symmetrical envelope, and require an iterative inference procedure, as described above for audition.

Temporal slowness constraints have also been used to model the spatiotemporal RFs of simple cells. The bubbles^26^ approach combines sparse and temporal coherence constraints with reconstruction. The resulting RFs show similar spatial and temporal properties to those found using ICA. A related framework is slow feature analysis (SFA)^7,37^, which enforces temporal smoothness by minimizing the derivative of unit responses over time, while maximising decorrelation between units. SFA has been used to model complex cell spatiotemporal RFs (over only two timesteps^7^), and a modified version has been used to model spatial (not spatiotemporal) RFs of simple cells^8^. These results are not directly comparable with our results or the spatiotemporal RFs of simple cells.

In the slowness framework, the features found are those that persist over time; the presence of such a feature in the recent past predicts that the same feature will be present in the near future. This is also the case for our predictive approach, which, additionally, can capture features in the past that predict features in the future that are subtly or radically different from themselves. The temporal prediction principle will also give different weighting to other features, as it values predictive capacity rather than temporal slowness^38^. In addition, although slowness models can be extended to model RFs over more than one timestep^7,22,26^, capturing temporal structure, they do not inherently give more weighting to information in the most recent past and therefore do not give rise to asymmetric temporal profiles in RFs.

There is one study that has directly examined temporal prediction as an objective for visual RFs in a manner similar to ours^39^. Here, as in our model, a single hidden layer feedforward neural network was used to predict the immediate future frame of a movie patch from its past frames. However, only two frames of the past were used in this study, so a detailed exploration of the temporal profile of the spatiotemporal RFs was not possible. Nevertheless, some similarities and differences in the spatial RFs between the two frames were noted, and some units had oriented RFs. In contrast to our model, however, many RFs were noisy and did not resemble those of simple cells. Potential reasons for this difference include the use of *L*_2_ rather than *L*_1_ regularization on the weights, an output nonlinearity not present in our model, the optimization algorithm used, network size, or the dataset.

Finally, it is worth noting the difference between our approach, which is concerned with temporal prediction, and another normative approach, known as the predictive coding framework^35,40,41^. The predictive coding framework has been used to explain both spatial RFs and higher order non-linear properties in visual cortex, but not the temporal properties of V1 RFs. Predictive coding generates top-down predictions of the current input it receives from lower levels, cancelling out or reducing activity in lower layers that is predictable by higher layers. The ‘prediction’ here does not refer to expectation of the future, but rather the output of the generative model for static input. In contrast, our model explicitly generates a prediction of future input that will be received by the model. Although predictive coding aims to efficiently represent all features of the current input, temporal prediction only aims to represent those features of current input that are predictive of the future^33,34^.

### Strengths and limitations of the temporal prediction model

Temporal prediction has several strengths as an objective function for sensory processing. First, it can capture underlying features in the world^1^; this is also the case with sparseness^4,5^ and slowness^37^, but temporal prediction will prioritise different features. Second, it can predict future inputs, which is very important for guiding action, especially given internal processing delays. Third, objectives such as efficient or sparse reconstruction retain everything about the stimulus, whereas an important part of neural information processing is the selective elimination of irrelevant information^42^. Prediction provides a good initial criterion for eliminating potentially unwanted information. Fourth, prediction provides a natural method to determine the hyperparameters of the model (such as regularization strength, number of hidden units, activation function and temporal window size). Other models select their hyperparameters depending on what best reproduces the neural data, whereas we have an independent criterion – the capacity of the network to predict the future. One notable hyperparameter is how many time-steps of past input to encode. As described above, this is naturally decided by our model because only time-steps that help predict the future have significant weighting. Fifth, the temporal prediction model computes neuronal activity without needing to settle to a steady state, unlike some other models^4–6,10,19,23,27,28^. For dynamic stimuli, a model that requires settling may not reach equilibrium in time to be useful. Sixth, and most importantly, temporal prediction successfully models many aspects of the RFs of primary cortical neurons. In addition to accounting for spatial and spectral tuning in V1 and A1, respectively, at least as well as other normative models, it reproduces the temporal properties of RFs, particularly the asymmetry of the envelopes of RFs, something few previous models have attempted to explain.

Although the temporal prediction model’s ability to describe neuronal RFs is high, the match with real neurons is not perfect. For example, the span of frequency tuning of our modelled auditory RFs is narrower than in A1 (Fig 6g-h). We also found an overrepresentation of vertical and horizontal orientations compared to real V1 data (Fig 7b). Some of these differences could be a consequence of the data used to train the model. Although the statistics of natural stimuli are broadly conserved^43^, there is still variation^44^, and the dataset used to train the network may not match the sensory world of the animal experienced during development and over the course of evolution. In future work, it would be valuable to explore the influence of natural datasets with different statistics, and also to match those datasets more precisely to the evolutionary context and individual experience of the animals examined. Furthermore, a comparison of the model with neural data from different species, at different ages, and reared in different environments would be useful.

Another cause of differences between the model and neural RFs may be the recording location of the RFs and how they are characterised. We used the primary sensory cortices as regions for comparison, because we performed transformations on the input data that are similar to the preprocessing that takes place in afferent subcortical structures. We spatially filtered the visual data in a similar way to the retina^4,5^, and spectrally decomposed the auditory data as in the inner ear, and then used time bins (5ms) which are coarser than, but close to, the maximum amplitude modulation period that can be tracked by auditory midbrain neurons^45^. However, primary cortex is not a homogenous structure, with neurons in different layers displaying certain differences in their response properties^46^. Furthermore, the methods by which neurons are sampled from the cortex may not provide a representative sample. For example, multi-electrode arrays tend to favour larger and more active neurons. In addition, the method and stimuli used to construct RFs from the data can bias their structure somewhat^21^.

The model presented here is based on a simple feedforward network with one layer of hidden units. This limits its ability to predict features of the future input, and to account for RFs with nonlinear tuning. More complex networks, with additional layers or recurrency may allow the model to account for more complex tuning properties, including those found beyond the primary sensory cortices. Careful, principled adjustment of the preprocessing, or different regularisation methods (such as sparseness or slowness applied to the units’ activities), may also help. Finally, the current model may eliminate some information that is useful for prediction of higher order statistical properties of the future input. Thus, the model may be improved further by adding feedback from higher areas or by adding an objective^3–5,31^ to reconstruct the past or present in addition to predicting the future.

To determine whether the model could help explain neuronal responses in higher areas, it would be useful to develop a hierarchical version of the temporal prediction model, applying the same model again to the activity of the hidden units rather than to the input. Another useful extension would be to see if the features learnt by the temporal prediction model could be used to accelerate learning of useful tasks such as speech or object recognition, by providing input or initialisation for a supervised or reinforcement learning network. Indeed, temporal predictive principles have been shown to be useful for unsupervised training of networks used in visual object recognition^47–50^.

Finally, a more detailed biological basis for our model would be valuable. In this study, we are agnostic as to the exact mechanism by which the RFs predicted by our model could be instantiated in the brain. The RFs could be hard-wired by evolution, given a species’ particular sensory environment, or they could be shaped during development by a predictive learning mechanism that may (or may not) continue to operate in adulthood. If RFs are acquired or refined through experience rather than simply hard-wired, then we would expect the brain to contain mechanisms capable of representing the accuracy of ongoing predictions about incoming sensory scenes; indeed, signals relating to prediction error have been found in A1^51^.

## Conclusion

We have shown that a simple principle - predicting the imminent future of a sensory scene from its recent past - explains many features of the RFs of neurons in both primary visual and auditory cortex. This principle may also account for neural tuning in other sensory systems, and may prove useful for the study of higher sensory processing and aspects of neural development and learning. While the importance of temporal prediction is increasingly widely recognized, it is perhaps surprising nonetheless that many basic tuning properties of sensory neurons, which we have known about for decades, appear, in fact, to be a direct consequence of the brain’s need to predict what will happen next.

## Methods

### Data used for model training and testing

#### Visual inputs

Videos (without sound, sampled at 25 fps) of wildlife in natural settings were used to create visual stimuli for training the artificial neural network. The videos were obtained from http://www.arkive.org/species, contributed by: BBC Natural History Unit, http://www.gettyimages.co.uk/footage/bbcmotiongallery; BBC Natural History Unit & Discovery Communications Inc., http://www.bbcmotiongallery.com; Granada Wild, http://www.itnsource.com; Mark Deeble & Victoria Stone Flat Dog Productions Ltd., http://www.deeblestone.com; Getty Images, http://www.gettyimages.com; National Geographic Digital Motion, http://www.ngdigitalmotion.com. The longest dimension of each video frame was clipped to form a square image. Each frame was then band-pass filtered^5^ and downsampled (using bilinear interpolation) over space, to provide 180x180 pixel frames. Non-overlapping patches of 20×20 pixels were selected from a fixed region in the centre of the frames, where there tended to be visual motion. The video patches were cut into sequential overlapping clips each of 8 frames duration. Thus, each training example (clip) was made up of a 20×20 pixel section of the video with a duration of 8 frames (320ms), providing a training set of *N*= ~500,000 clips from around 5.5 h of video, and a validation set of *N*= ~100,000 clips. Finally, the training and validation sets were normalised by subtracting the mean and dividing by the standard deviation (over all pixels, frames and clips in the training set). The goal of the neural network was to predict the final frame (the ‘future’) of each clip from the first 7 frames (the ‘past’).

#### Auditory inputs

Auditory stimuli were compiled from databases of human speech (~60%), animal vocalizations (~20%) and sounds from inanimate objects found in natural settings (e.g. running water, rustling leaves; ~20%). Stimuli were recorded using a Zoom H4 or collected from online sources. Natural sounds were obtained from www.freesound.org, contributed by users sedi, higginsdj, jult, kvgarlic, xenognosis, zabuhailo, funnyman374, videog, j-zazvurek, samueljustice00, gfrog, ikbenraar, felix-blume, orbitalchiller, saint-sinner, carlvus, kyster, vflefevre, hitrison, willstepp, timbahrij, xdimebagx, r-nd0mm3m, the-yura, rsilveira-88, stomachache, foongaz, edufigg, yurkobb, sandermotions, darius-kedros, freesoundjon-01, dwightsabeast, borralbi, acclivity, J.Zazvurek, Zabuhailo, soundmary, Darius Kedros, Kyster, urupin, RSilveira and freelibras. Human speech sounds were obtained from http://databases.forensic-voice-comparison.net/^52,53^.

Each sound was sampled at (or resampled to) 44.1 kHz and converted into a simple ‘cochleagram’, to make it more analogous to the activity pattern that would be passed to the auditory pathway after processing by the cochlea. To calculate the cochleagram, a power spectrogram was computed using 10ms Hanning windows, overlapping by 5ms (giving time steps of 5ms). The power across neighbouring Fourier frequency components was then aggregated into 32 frequency channels using triangular windows with a base width of 1/3 octave whose centre frequencies ranged from 500 to 17,827Hz (1/6^th^ octave spacing, using code adapted from melbank.m, http://www.ee.ic.ac.uk/hp/staff/dmb/voicebox/voicebox.html). The cochleagrams were then decomposed into sequential overlapping clips, each of 43 time steps (415 ms) in duration, providing a training set of ~1,000,000 clips (~1.3 hours of audio) and a validation set of ~200,000 clips. To approximately model the intensity compression seen in the auditory nerve^54^, each frequency band in the stimulus set was divided by the median value in that frequency band over the training set, and passed through a hill function, defined as *h(x) = cx*/(*1* + *cx*) with *c*=0.02. Finally, the training and cross-validation sets were normalised by subtracting the mean and dividing by the standard deviation over all time steps, frequency bands and clips in the training set. The first 40 time steps (200 ms) of each clip (the ‘past’) were used as inputs to the neural network, whose aim was to predict the content (the ‘future’) of the remaining 3 bins (15 ms).

#### Addition of Gaussian noise

To replicate the effect of noise found in the nervous system, Gaussian noise was added to both the auditory and visual inputs with a signal-to-noise ratio (SNR) of 4. While the addition of noise did not make substantial differences to the RFs of units trained on visual inputs, this improved the similarity to the data when the model was trained on auditory inputs. The results from training the network on inputs without added noise are shown for auditory inputs in Fig. 4-Fig. Supplement 5 and Fig. 6-Fig. Supplement 1 and for visual inputs in Fig. 4-Fig. Supplement 6-7 and Fig. 7-Fig. Supplement 2. The results from the sparse coding model were similar in both cases for inputs with and without noise (Figs. 5-6, Fig. 5-Fig. Supplements 1-5, Fig. 6-Fig. Supplement 1, Fig. 7-Fig. Supplements 1,3).

### Temporal prediction model

#### The model and cost function

The temporal prediction model was implemented using a standard fully connected feed-forward neural network with one hidden layer. Each hidden unit in the network computed the sum of linearly weighted inputs, and its output was determined by passing this sum through a monotonic nonlinearity. This nonlinearity *s* = f(*a*) was either a logistic function f(*a*) = 1/(1+exp(*-a*)) or a similar nonlinear function (such as tanh). For results reported here, we used the logistic function, though obtained similar results when we trained the model using f(*a*) = tanh(*a*). For comparison, we also trained the model replacing the nonlinearity with a linear function, where f(*a*) =*a*. In this case, we found that the RFs tended to be punctate in space or frequency and did not typically show the alternating excitation and inhibition characteristic real neurons in A1 and V1.

Formally, for a network with *i* = 1 to *I* input variables, *k* = 1 to *K* output units and a single layer of *j* = 1 to *J* hidden units, the output *s_jn_* of hidden unit *j* for clip *n* is given by:

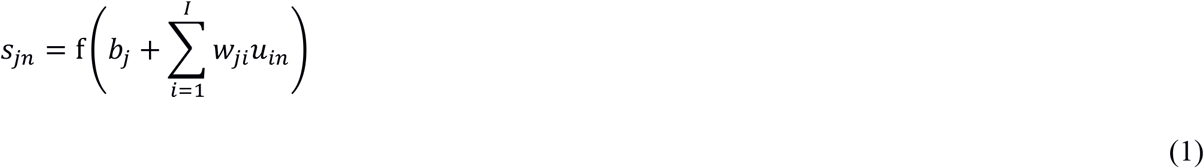

The value *u_in_* of input variable *i* for clip *n* is simply the value for a particular pixel and frame of the ‘past’ in preprocessed visual clip *n* (*I* = 20×20×7 = 2800), or the value for a particular frequency band and time step of the ‘past’ of cochleagram clip *n* (*I* = 32×40 = 1280). The subscript *n* has been dropped for clarity in the figures (Fig. 1). The parameters in Equation 1 are the connective input weights *w_ji_* (between each input variable *i* and hidden unit *j*), and the bias *b_j_* (of hidden unit *j*).

The activity 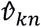 of each output unit *k*, which is the estimate of the true future *v_kn_* given the past *u_in_*, is given by:

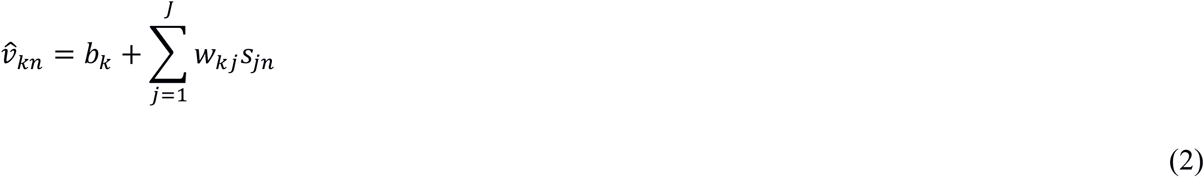

The parameters in Equation 2 are the connective output weights *w_kj_* (between each hidden unit *j* and output unit *k*) and the bias *b_k_* (of output unit *k*). The activity 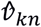 of output unit *k* for clip *n* is the estimate for a particular pixel of the ‘future’ in the visual case (*K* = 20×20×1 = 400), or the value for a particular frequency band and time step of the ‘future’ in the auditory case (*K* = 32×3 = 96).

The parameters *w_ji_*, *w_kj_*, *b_j_*, and *b_k_* were optimised for the training set by minimizing the cost function given by:

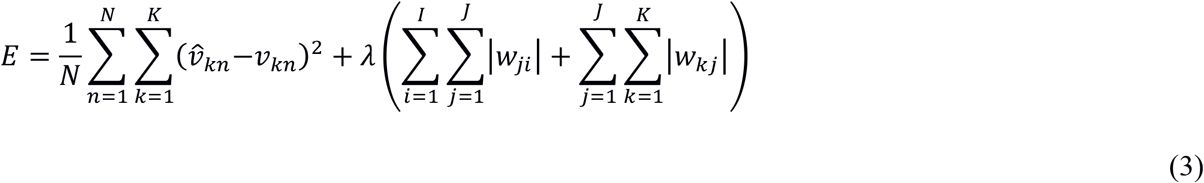

Thus, *E* is the sum of the squared error between the prediction 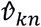 and the target *v_kn_* over all *N* training examples, plus an *L*_1_ regularisation term, which is proportional to the sum of absolute values of all weights in the network and its strength is determined by the hyper-parameter *λ*. This regularisation tends to drive redundant weights to near zero and provides a parsimonious network.

#### Implementation details

The networks were implemented in Python (https://lasagne.readthedocs.io/en/latest/;http://deeplearning.net/software/theano/). The objective function was minimised using backpropagation as performed by the Adam optimisation method^55^. An alternative implementation of the model was also made in MATLAB using the Sum-of-Functions Optimiser^56^ (https://github.com/Sohl-Dickstein/Sum-of-Functions-Optimizer) to train the network using backpropagation. Training examples were split into minibatches of approximately 7000 training examples each.

During model network training, several hyperparameters were varied, including the regularisation strength (λ), the number of units in the hidden layer and the nonlinearity used by each hidden unit. For each hyperparameter setting, the training algorithm was run for 1000 iterations. Running the network for longer (10000 iterations) showed negligible improvement to the prediction error (as measured on the validation set) or change in RF structure.

The effect of varying the number of hidden units and λ on the prediction error for the validation set is shown in Fig. 8. In both the visual and auditory case, the results presented are the networks that predicted best on the validation set after 1000 iterations through the training data. For the auditory case, the settings that resulted in the best prediction were 1600 hidden units and λ = 10^−3.5^, while in the visual case, the optimal settings were 1600 hidden units and *λ* = 10^−3.75^.

#### Model receptive fields

In the model, the combination of linear weights and nonlinear activation function are similar to the basic linear non-linear (LN) model^57–61^ commonly used to describe neural RFs. Hence, the input weights between the input layer and a hidden unit of the model network are taken directly to represent the unit’s RF, indicating the features of the input that are important to that unit.

Because of the symmetric nature of the sigmoid function, f(*a*) = 1-f(-*a*), with appropriate modification of the biases, a hidden unit has the same influence on the prediction if its input and output matrices are both multiplied by −1. That is, for unit *j*, if we convert w*_ij_* to −w*_ij_*, w*_jk_* to −w*_jk_*, b*_j_* to −b*_j_*, and b*_k_* to −b*_k_* + w*_jk_*, this will have no effect on the prediction or the cost function. This can be done independently for each hidden unit. Hence, the sign of each unit’s RF could equally be positive or negative and have the same result on the predictions given by the network. However, we know that auditory units always have leading excitation (Fig. 3). Hence, for both the predictive model and for the sparse coding model, we assume leading excitation for each unit. This was done for all analyses.

As more units are added to the model network, the number of inactive units increases. To account for this, we measured the relative strength of all input connections to each hidden unit by summing the square of all input weights for that unit. Units for which the sum of square input weights was <1% of the maximum strength for the population were deemed to be inactive and excluded from all subsequent analyses. The difference in connection strength between active and inactive units was very distinct; a threshold <0.0001% only marginally increases the number of active units.

### Sparse coding model

The sparse coding model was used as a control for both visual and auditory cases. The Python implementation of this model (https://github.com/zayd/sparsenet)was trained using the same visual and auditory inputs used to train the predictive model. The training data were divided into minibatches which were shuffled and the model optimised for one full pass through the data. Inference was performed using the Fast Iterative Shrinkage and Thresholding (FISTA) algorithm. A sparse *L*_1_ prior with strength λ, was applied to the hidden unit activities. A range of λ-values and hidden unit numbers were tried (Fig. 8-Fig. Supplement 1). The learning rate and batch size were also varied until reasonable values were found. As there was no independent criterion by which to determine the ‘best’ settings, we chose the network that produced basis functions that were deemed, by inspection, to be the most similar to A1 or V1 RFs. In this manner, we selected a sparse coding network with 800 hidden units, λ=10^0.5^, learning rate = 0.01 and 100 mini-batches in the auditory case (Figs. 5-6). In the visual case, the network selected had the same settings except for the number of hidden units, which was 400 (Fig. 5-Fig. Supplements 1-2 and Fig. 7-Fig. Supplement 1). Although these sparse coding basis functions are projective fields, they tend to be similar in structure to receptive fields^4,5^, and, for simplicity, are referred to as RFs.

### Auditory receptive field analysis

#### In vivo A1 RF data

Auditory RFs of neurons in the primary auditory cortical fields A1 and AAF of 5 anesthetised ferrets were recorded and used as a basis of comparison for the RFs of model units trained on auditory stimuli. Systematic differences in response properties of A1 and AAF neurons are minor and not relevant for this study, and for simplicity here, we refer to neurons from either primary field indiscriminately as “A1 neurons”. The neural responses were recorded using multi-electrode arrays (Neuronexus), the methods are detailed in an earlier study^21^, and described briefly below. The neuronal recordings used the “BigNat”stimulus set^21^, which consists of natural sounds including animal vocalizations (e.g., ferrets and birds), environmental sounds (e.g., water and wind), and speech. To identify those neural units that were driven by the stimuli, we calculated a “noise ratio” statistic^61,62^ for each units and excluded from further analysis any neurons with a noise ratio >40. In total, driven spiking responses of 114 units (75 single unit, 39 multi-unit) were recorded to this stimulus set. Then, the auditory (spectrotemporal) RF of each unit was constructed using a previously described method^21^. Briefly, linear regression was performed in order to minimise the squared error between each neuron’s spiking response over time and the cochleagram of the stimuli that gave rise to that response. The method used was exactly the same as in our earlier study^21^, except that *L*_1_ rather than *L*_2_ regularisation was used to constrain the regression. The spectrotemporal RFs of these neurons took the same form as the inputs to the model neural network (i.e., 32 frequencies and 40 time-steps over the same range of values) and were therefore comparable to the model units’ RFs. In order to account for the latency of auditory cortical responses, the first two time-steps (10ms) of the neuronal responses were removed, leaving 38 time-steps.

#### Multi-Dimensional Scaling (MDS)

To get a non-parametric indication of how similar the model units’ RFs were to those of real A1 neurons, each RF was embedded into a 2-dimensional space using MDS (Fig. 6a and Fig. 6-Fig. Supplement 1a). First, 100 units each from the temporal prediction and sparse coding models and from the real population were chosen at random. To ensure that the model RFs were of the same dimensionality prior to embedding, the final two timesteps of each model RF were removed.

#### Measuring temporal and frequency spans of RFs

We quantified the span, over time and frequency, of the excitatory and inhibitory subfields of each RF. To do this, each RF was first separated into excitatory and inhibitory subfields, where the excitatory subfield was the RF with negative values set to 0, and the inhibitory subfield the RF with positive values set to 0. In some cases, model units did not exhibit notable inhibitory subfields. To account for this, the power contained in each subfield was calculated (sum of the squares of the subfield). Inhibitory subfields with < 5% of the power of that unit’s excitatory subfield were excluded from further analysis. According to this criterion, 42 of 167 active units in the temporal prediction model and 137 of 800 units in the sparse model did not display inhibition.

Singular value decomposition (SVD) was performed on each subfield separately, and the first pair of singular vectors was taken, one of which is over time, the other over frequency. For the excitatory subfield, the temporal span was measured as the proportion of values in the temporal singular vector that exceeded 50% of the maximum value in the vector. The same analysis provided the temporal span for the inhibitory subfield. Similarly, we measured the frequency spans of the RFs by applying this measure to the frequency singular vectors of the excitatory and the inhibitory subfields.

We also examined, for both real and model RFs, the mean power for each of the 38 time steps in the RFs (Fig. 6b), which was calculated as the mean of the squared RF values, over all frequencies and RFs, at each time step.

#### Mean KS measure

To compare each network’s units with those recorded in A1 (Fig. 3), the two-sample Kolmogorov-Smirnov (KS) distance between the real and model distribution was measured for both the temporal and spectral span of the excitatory and inhibitory subfields (e.g. the distributions in Fig. 6d-e and Fig. 6g-h). These four KS measures were then averaged to give a single mean KS measure for each network, indicating how closely the temporal and frequency characteristics of real and model units matched on average for that network. The KS measure is low for similar distributions and high for distributions that diverge greatly. Thus networks whose units display temporal and frequency tuning characteristics that match those of real neurons more closely give rise to a lower mean KS measure.

### Visual receptive field analysis

#### In vivo V1 RF data

Visual RFs measured using recordings from V1 simple cells were compared against the model (Fig. 2c, and Fig. 7a (cat^24^) and Fig. 7c (cat^15^, mouse^30^ and monkey^17^)). The in vivo data were taken from the authors’ website^16^ or extracted from relevant papers^15^ or provided by the authors^24,30^.

#### Fitting Gabors

In order to quantify tuning properties of the model’s visual RFs, 2D Gabors were fitted to the optimal time-step of each unit’s response^15,17^. This allowed comparison to previous experimental studies which parameterised real RFs by the same method^17^. The optimal time-step was defined^17^ as the time-step of the unit’s response which contained the most power (sum of square values). The Gabor function has been shown to provide a good approximation for most spatial aspects of simple visual RFs^15,17^. The 2D Gabor is given as:

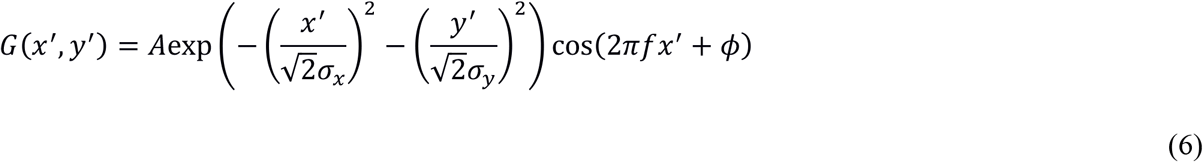

where, the spatial coordinates (*x*’, *y*’) are acquired by translating the centre of the RF (*x*_0_, *y*_0_) to the origin and rotating the RF by its spatial orientation *θ*:

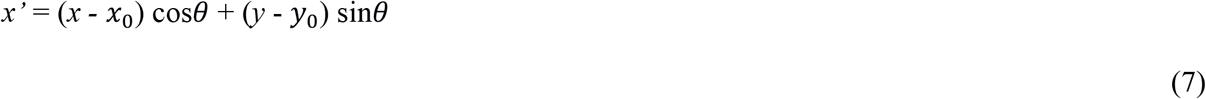

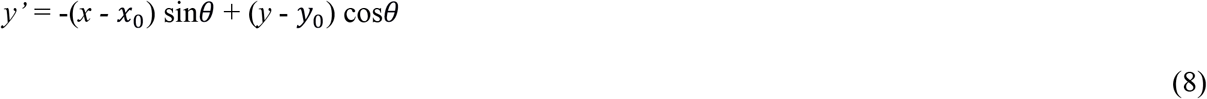

*σ_x_* and *σ_y_* provide the width of the Gaussian envelope in the *x*’ and *y*’ directions, while *f* and *ϕ* parameterise the spatial frequency and phase of the sinusoid along the *x*’ axis. *A* parameterises the height of the Gaussian envelope.

For each RF, the parameters (*x*_0_, *y*_0_, *σ_x_*, *σ_y_*, *θ*, *f*, *ϕ*) of the Gabor were fitted by minimizing the mean squared error between the Gabor model and the RF using the minFunc minimization package (http://www.cs.ubc.ca/~schmidtm/Software/minFunc.html). In order to avoid local minima, the fitting was performed in two steps. First, the spatial RF was converted to the spectral domain using a 2D Fourier transform. Since the Fourier transform of a 2D Gabor is a 2D Gaussian^15^, which is easier to fit, an estimate of many of the parameters was obtained by first fitting a 2D Gaussian in the spectral domain. Using the parameters obtained from the spectral fitting as initial estimates, a 2D Gabor was then fitted to the original RF in the spatial domain. The fitted parameters provided a good estimate of the units’ responses, with residual errors between the spatial responses and the corresponding Gabor fits being small and lacking spatial structure, and the median pixel-wise correlation coefficient of the Gabor fits for the temporal prediction model units was 0.96. Units whose fitted Gabors had a poor fit (those with a correlation coefficient <0.7; 14 units) were excluded from further analysis. We also excluded units with a high correlation coefficient (>0.7) if the centre position of the Gabor was estimated to be outside the RF, and hence only the Gabor’s tail was being fitted to the response (91 units), and those for which the estimated standard deviation of the Gaussian envelope in either x or y was <0.5 pixels, which meant very few non-negligible pixel values were used to constrain the parameters (267 units). Together, these exclusion criteria (which sometimes overlapped), led to 395 of the 1493 responsive units being excluded for the temporal prediction model.

#### 2D spatiotemporal receptive fields

In order to better view their temporal characteristics we collapsed the 3D spatiotemporal real and model RFs (space-space-time) along a single spatial direction to create 2D spatiotemporal (spacetime) representations^16^. First, we determined the 3D RFs’ optimal time step (the time step with the largest sum of squared values). We then acquired the rotation and translation that centres the RF on zero and places the oriented bars parallel to the y-axis at the optimal time step from the Gabor parameterisation of each unit at its optimal time step. We applied this fixed transformation to each time step and collapsed the RF by summing the activity along the newly defined y-axis. The resulting 2D (space-time) RFs provide intuitive visualisation of the RF across time, while losing minimal information. For the RFs of real neurons^24^, the most recent time bin (40ms) of the 3D and 2D spatiotemporal RFs were removed to account for the latency of V1 neurons (Fig 2c; 7a).

#### Estimating space-time separability

The population of model units contained both space-time (ST) separable and inseparable units. First the two spatial dimensions of the 20x20x7 3D RF were collapsed to a single vector to yield a single 400x7 matrix. The SVD of this matrix was then taken and the singular values examined. If the ratio between the second and first singular value was ≥0.5, the unit was deemed to be inseparable. Otherwise, the unit was deemed to be separable. Examining the 20x7 2D spatiotemporal RFs (obtained as outlined in the preceding section; Fig. 4-Fig. Supplement 3) showed this to be an accurate way of separating space-time separable and inseparable units.

#### Spatial RF structure

For comparison with the real V1 RF and previous theoretical studies, the width and length of our model’s RFs were measured relative to their spatial frequency^17^. Here, *n_x_* = *σ_x_f* gives a measure of the length of the bars in the RF, while *n_y_* = *n_y_f* gives a measure of the number of oscillations of its sinusoidal component. Thus, in the *n*_*y*_, *n_x_* plane, blob-like RFs with few cycles lie close to the origin, while stretched RFs with many subfields lie away from the origin. RFs with values high along the *n_y_* axis, have many bars, while those far along the *n_x_* axis have long bars. As in Ringach^17^ only spacetime separable units were included in this analysis.

#### Temporal weighting profile of the population

The mean power for each of the 7 time steps of the RFs was examined for both real and model populations (Fig. 7a). The temporal weighting profile was calculated as the mean, over space and the population, of the squared values of the 2D spatiotemporal RFs at each time step.

#### Peak temporal frequency

The 2D spatiotemporal RFs were also useful for calculating further temporal response properties of the model. The temporal frequency was calculated as the peak temporal frequency of each spatiotemporal RF as measured from its 2D Fourier transform.

### Code and data availability

All code used in this study was implemented in MATLAB and Python. We would be happy to provide our code to reviewers and will upload our code to Github on acceptance. The raw auditory experimental data is available at https://osf.io/ayw2p/. The movies and sounds used for training the models are all publicly available at the websites detailed in the Methods.

## Acknowledgments

Nicol Harper was supported by a Sir Henry Wellcome Postdoctoral Fellowship (WT082692) and other Wellcome Trust funding (WT076508AIA, WT108369/Z/2015/Z), by the Department of Physiology, Anatomy and Genetics at the University of Oxford, by Action on Hearing Loss (PA07), and by the Biotechnology and Biological Sciences Research Council (BB/H008608/1). Yosef Singer and Yayoi Teramoto were supported by the Clarendon Fund. Yayoi Teramoto was supported by the Wellcome Trust (10525/Z/14/Z). Andrew King and Ben Willmore were supported by the Wellcome Trust (WT076508AIA, WT108369/Z/2015/Z). We thank Bruno Olshausen for discussions on his model.

## Author contributions

Conceived and designed study: NH. Wrote code for models: NH, YS, YT. Wrote code for data analysis: YS, NH, BW. Performed modelling experiments: YS. Performed physiological experiments: BW, NH. Supervised project: NH, BW, JS, AK. Wrote manuscript: YS, NH, BW, YT, AK, JS.

## Supplementary Materials

**Figure 4-Figure Supplement 1.**
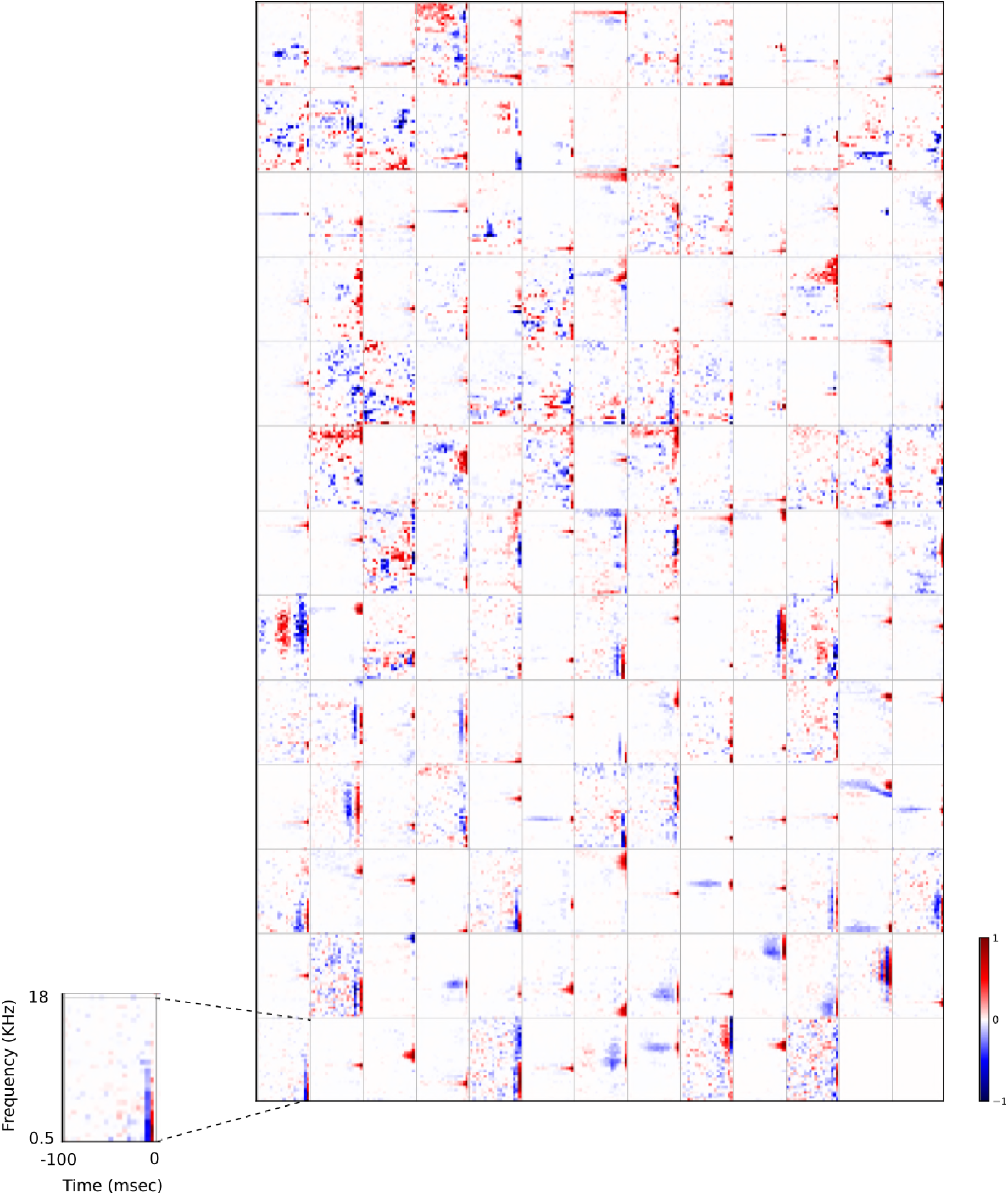
Full set of auditory RFs of the temporal prediction model units shown on a finer timescale. All details are as in Figure 3, but the only the most recent 100ms of the response profile is shown in order to illustrate details of the RFs. Inset shows axes.

**Figure 4-Figure Supplement 2.**
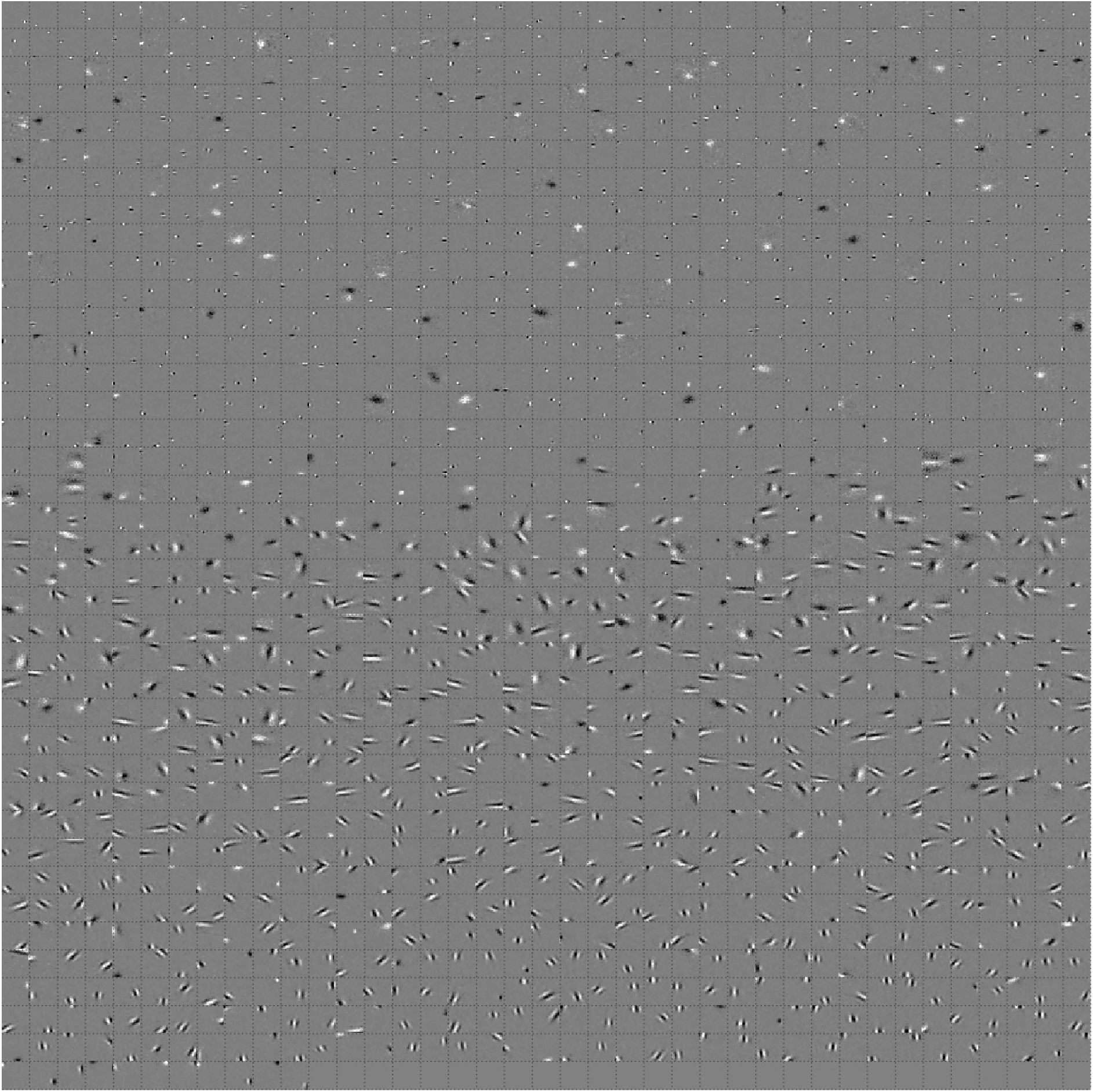
Full set of visual spatial RFs of the temporal prediction model units. Model units are the same as those used in Figure 7a. Example units in Fig. 2 come from this set. Each square represents the spatial RF of a single unit, shown at its best timestep. The best timestep was determined by selecting the timestep for which the power (sum of squares) of the RF was greatest. White – excitation, black - inhibition.

**Figure 4-Figure Supplement 3.**
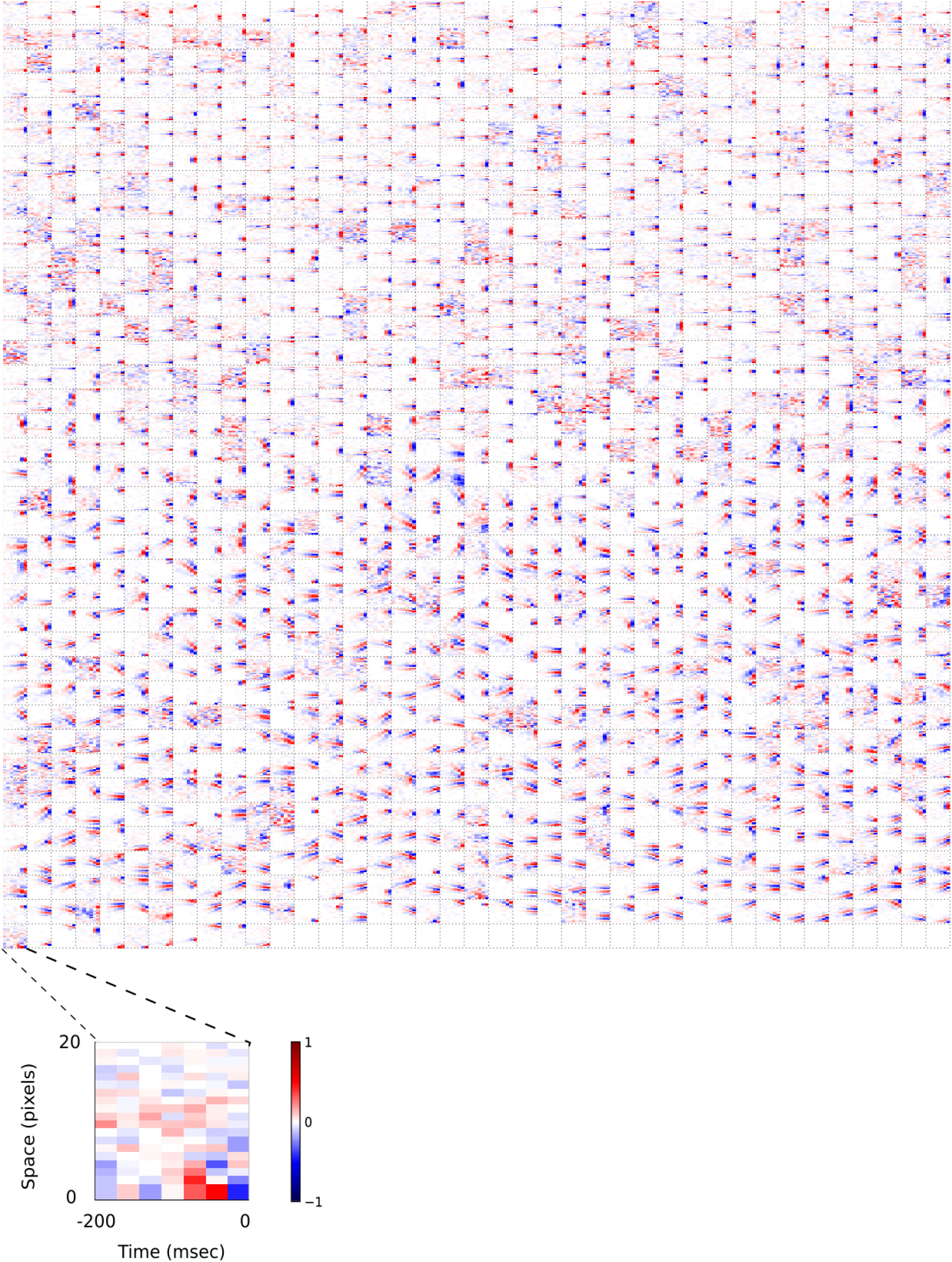
Visual 2D (space-time) spatiotemporal RFs of temporal prediction model units. Model units are the same as those used in Fig. 7. Each square represents the 2D spatiotemporal RF of a single unit corresponding to the unit in the same position in Fig. 4-Fig. Supplement 2, obtained by summing across space along the axis of the orientation for that unit. Red – excitation, blue - inhibition. Inset shows axes.

**Figure 4-Figure Supplement 4.**
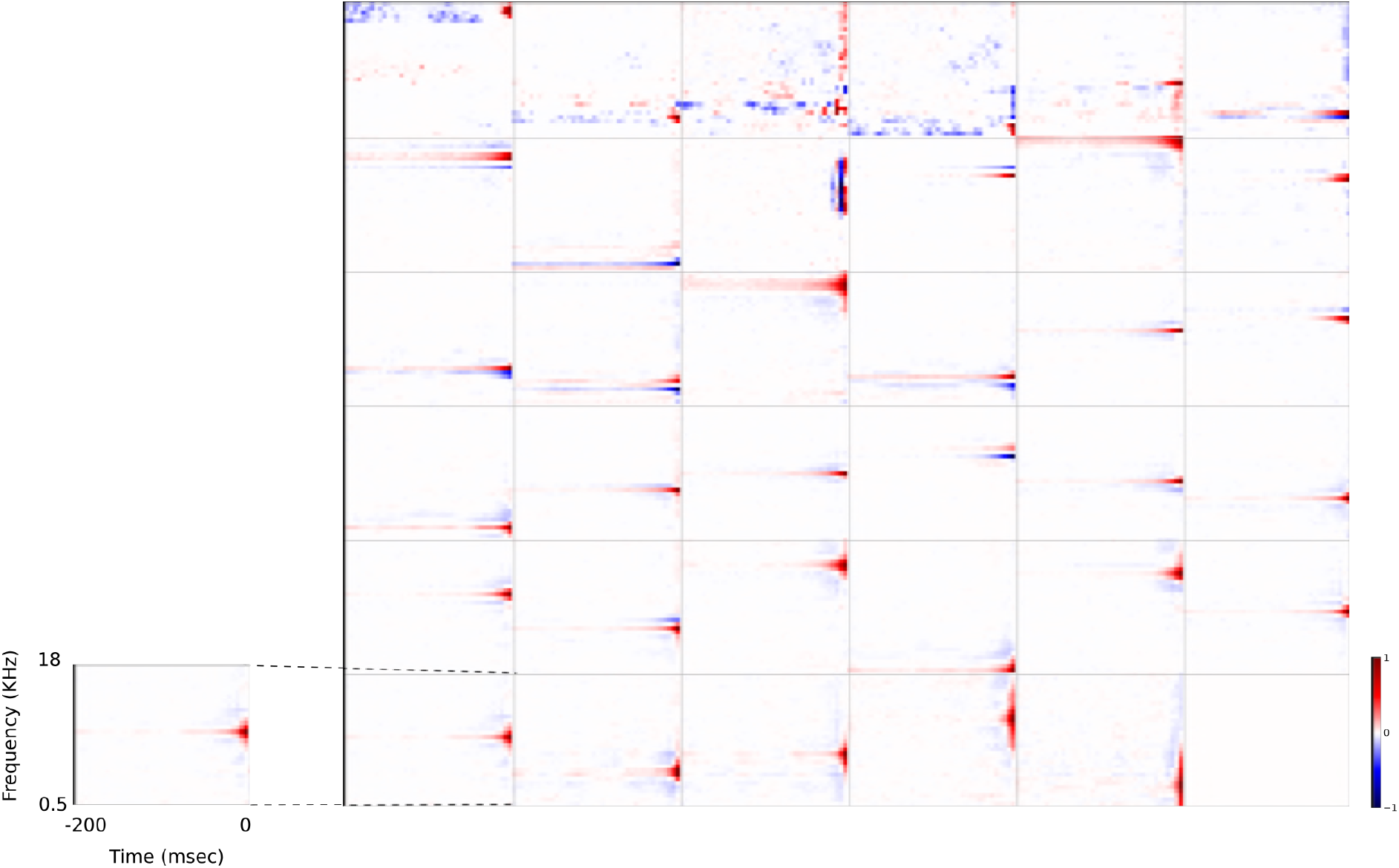
Full set of auditory RFs of the temporal prediction model units using a linear activation function. Units were obtained by training the model with 1600 hidden units on auditory inputs. The hidden unit number and L1 weight regularization strength (10^−3.25^) was chosen because they result in the lowest MSE on the prediction task, as measured using a cross validation set. Almost all hidden units’ weight matrices decayed to near zero during training (due to the L1 regularization), leaving 35 active units. Inactive units were excluded from analysis and are not shown. Red – excitation, blue - inhibition. Inset shows axes.

**Figure 4-Figure Supplement 5.**
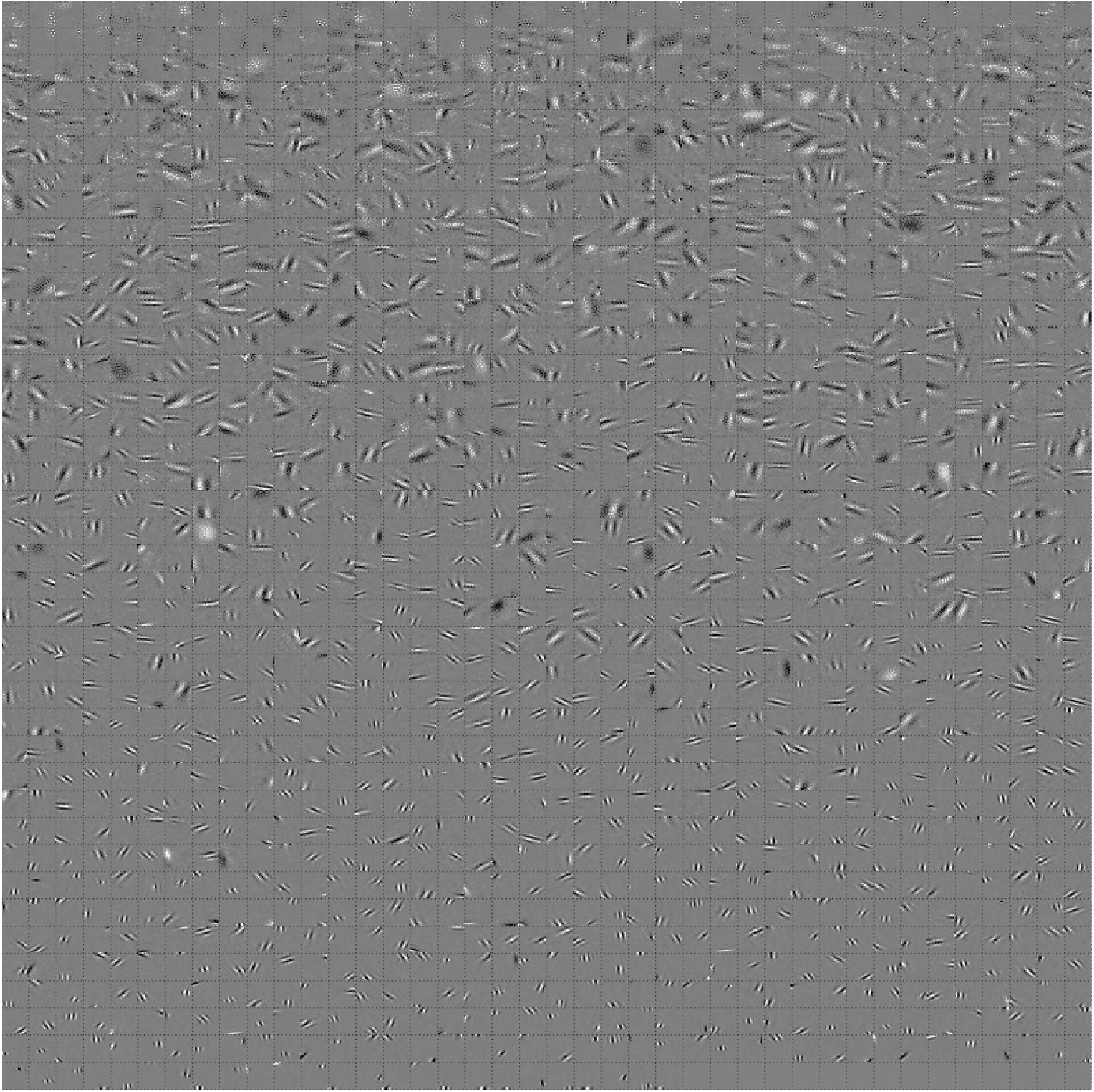
Full set of auditory RFs of the temporal prediction model units trained on auditory inputs without added noise. Units were obtained by training the model with 1600 hidden units on auditory inputs. The hidden unit number and L1 weight regularization strength (10^−4^) was chosen because it results in the lowest MSE on the prediction task, as measured using a cross validation set. Many hidden units’ weight matrices decayed to near zero during training (due to the L1 regularization), leaving 465 active units. Inactive units were excluded from analysis and are not shown. Red – excitation, blue - inhibition. Inset shows axes.

**Figure 4-Figure Supplement 6.**
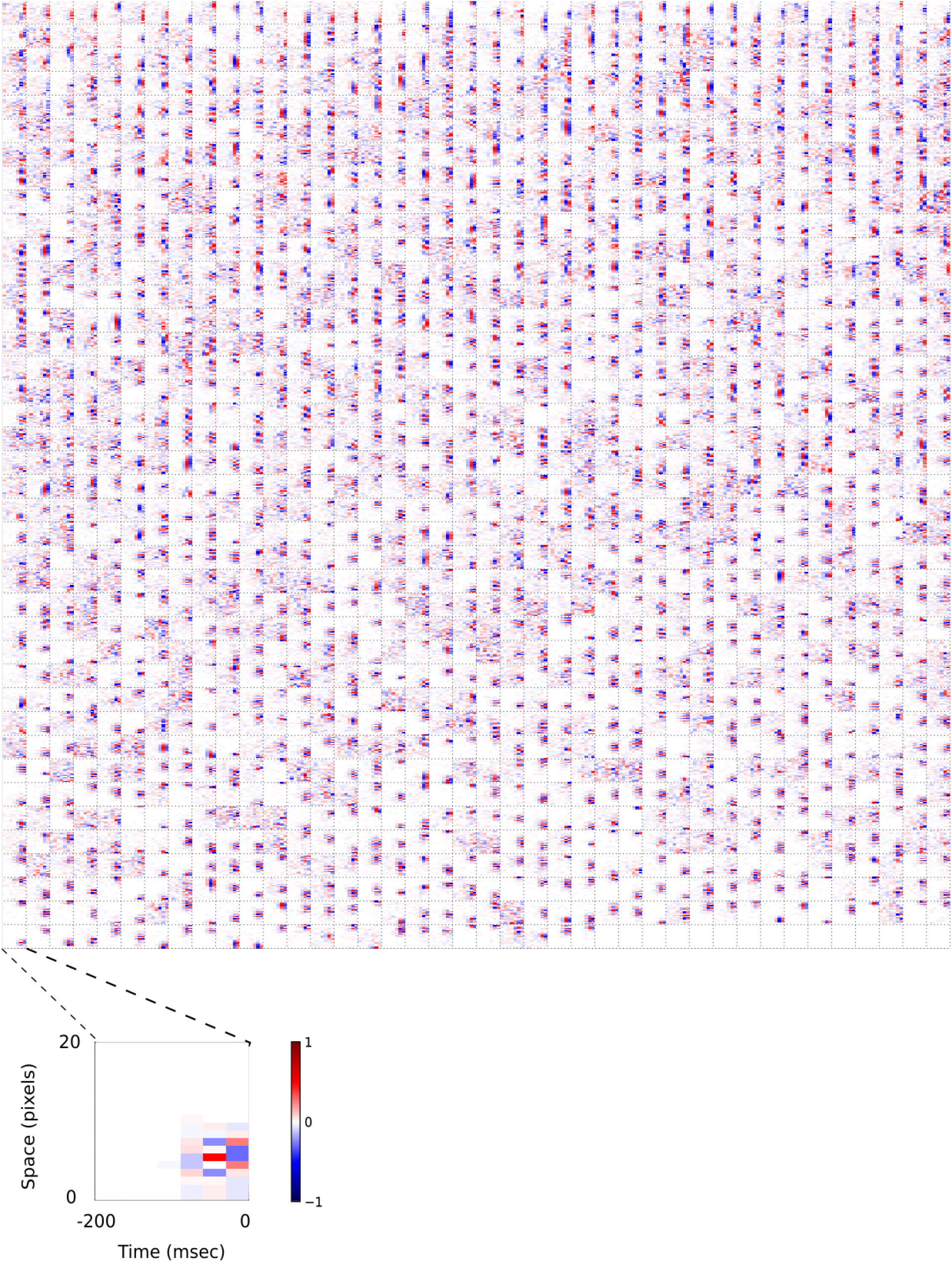
Full set of visual spatial RFs of temporal prediction model units trained on visual inputs without added noise. Model units were obtained by training the model with 1600 hidden units on visual inputs. The hidden unit number and L1 weight regularization strength (10^−4^) was chosen because it results in the lowest MSE on the prediction task, as measured using a cross validation set. Some hidden units’ weight matrices decayed to near zero during training (due to the L1 regularization), leaving 1585 active units, which were included in analysis. Inactive units were excluded from analysis. Each square represents the spatial RF of a single unit, shown at its best timestep. The best timestep was determined by selecting the timestep for which the power (sum of squares) of the RF was greatest. White – excitation, black - inhibition.

**Figure 4-Figure Supplement 7.**
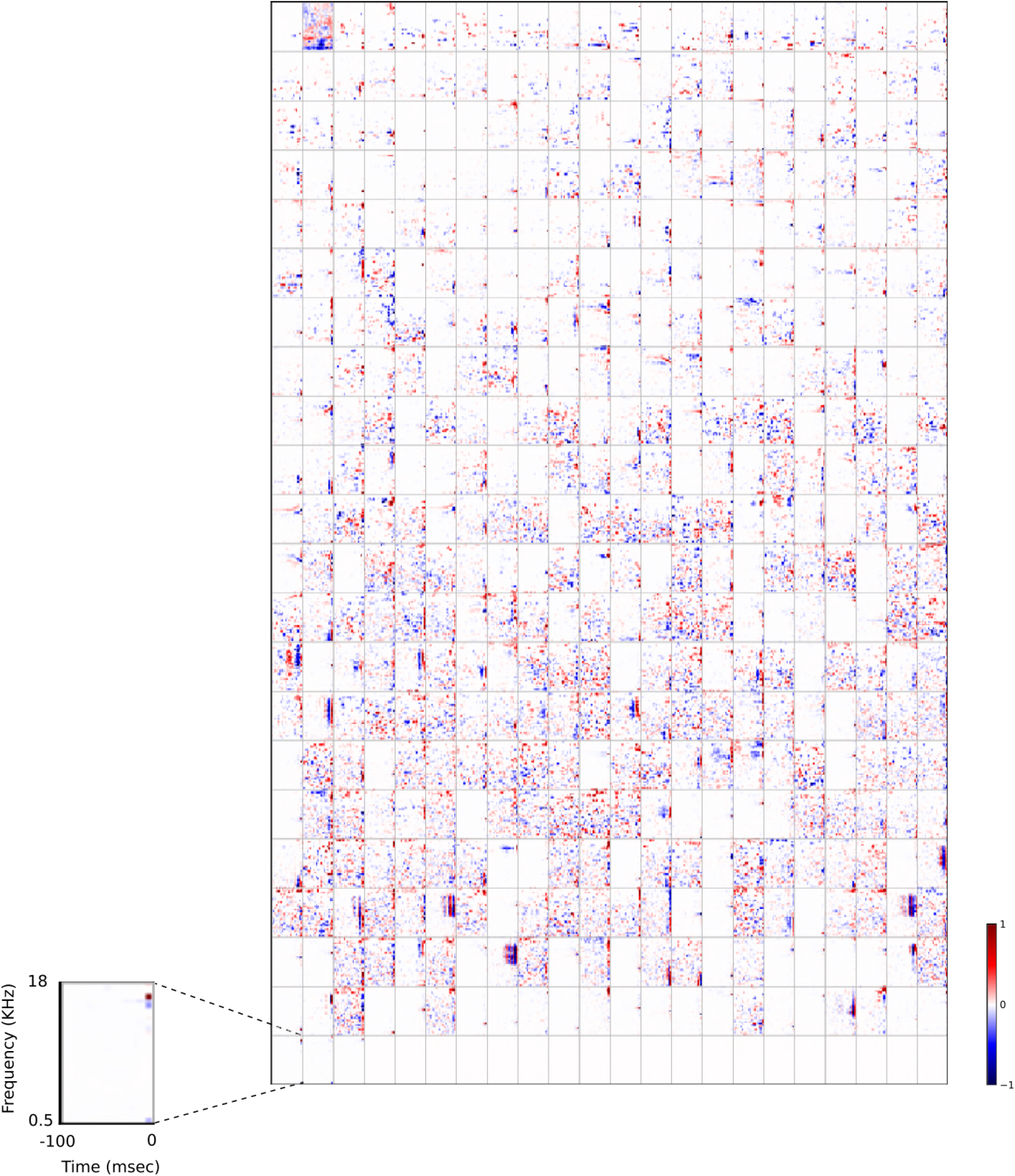
2D (space-time) visual spatiotemporal RFs of temporal prediction model units trained on visual inputs without added noise. Obtained from the same units shown in Fig. 4-Fig. Supplement 6 using methods outlined in Fig 2c. Red – excitation, blue - inhibition. Inset shows axes.

**Figure 5-Figure Supplement 1.**
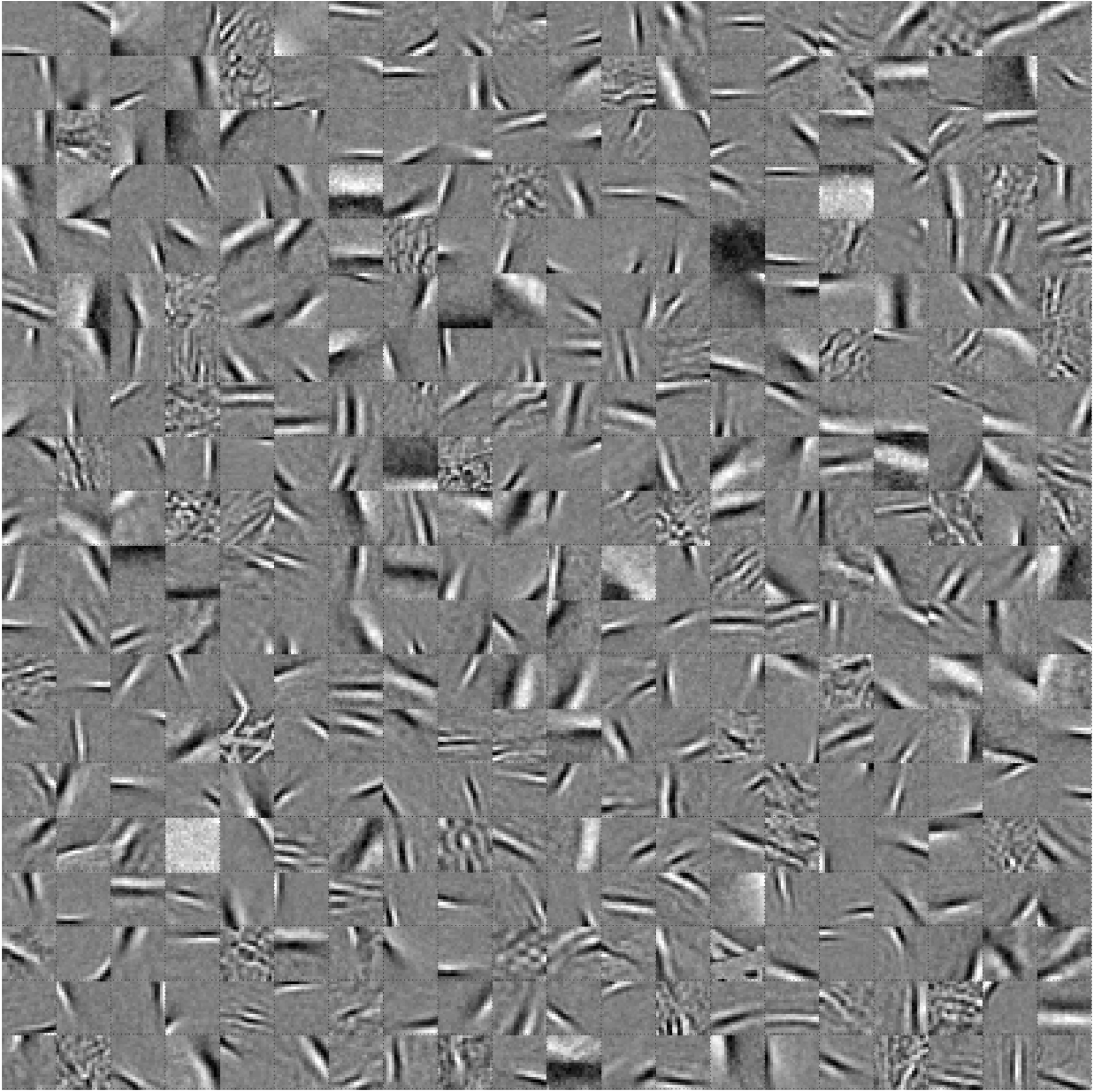
Full set of visual spatial RFs of sparse coding model units. Model units were obtained by training the sparse coding model with 400 hidden units on identical visual inputs used to train the temporal prediction model. The model configuration (400 hidden units, *L*_1_ sparsity strength of 10^0.5^ on the unit activities) was chosen because it resulted in the RFs that look most like the RFs of V1 simple cells as determined by visual inspection. Each square represents the spatial RF of a single unit, shown at its best timestep. The best timestep was determined by selecting the timestep for which the power (sum of squares) of the RF was greatest. White – excitation, black - inhibition.

**Figure 5-Figure Supplement 2.**
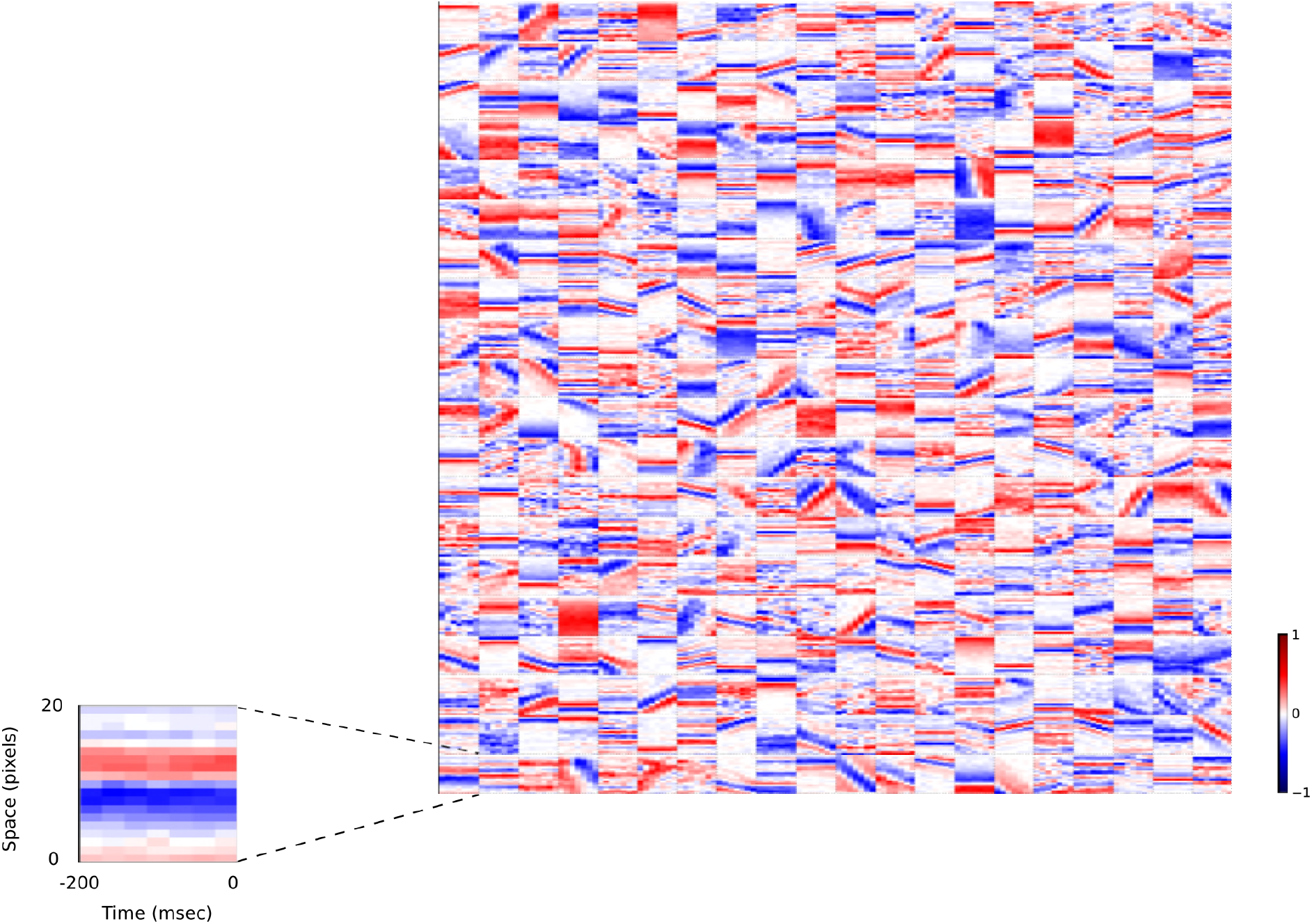
2D (space-time) visual spatiotemporal RFs of sparse coding model units. Obtained from the same units shown in Fig. 5-Fig. Supplement 1 using methods outlined in Fig 2c. Red – excitation, blue - inhibition. Inset shows axes.

**Figure 5-Figure Supplement 3.**
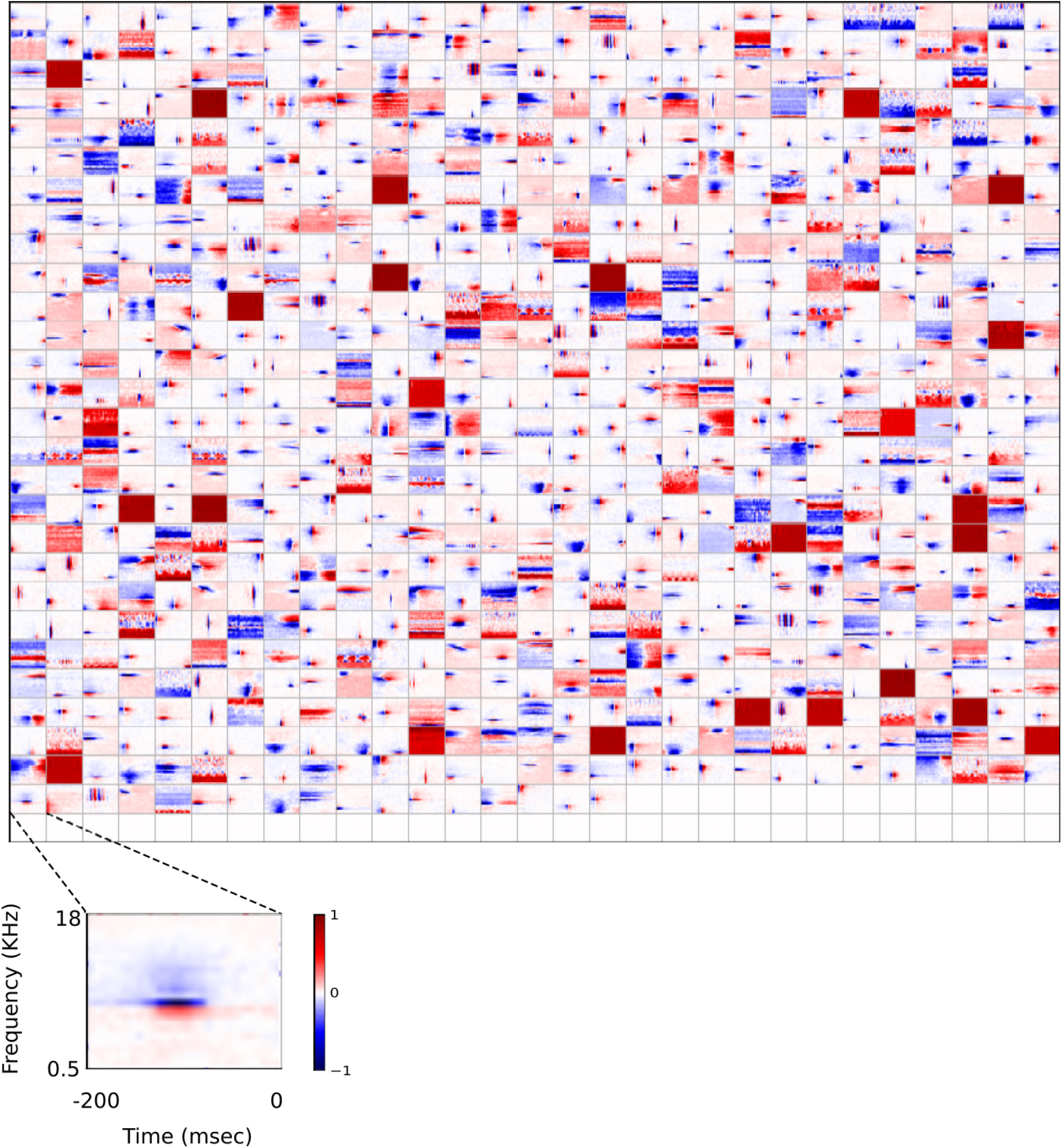
Full set of auditory RFs of sparse coding model trained on auditory inputs without added noise. Units were obtained by training the sparse coding model with 800 units on the identical auditory inputs used to train the network shown in Fig. 7-Fig. Supplement 2. L1 regularization of 10^0.05^ was applied to the hidden units’ activities. This network configuration was selected as it produced unit RFs that most closely resembled those recorded in A1, as determined by visual inspection. Red – excitation, blue - inhibition. Inset shows axes.

**Figure 5-Figure Supplement 4.**
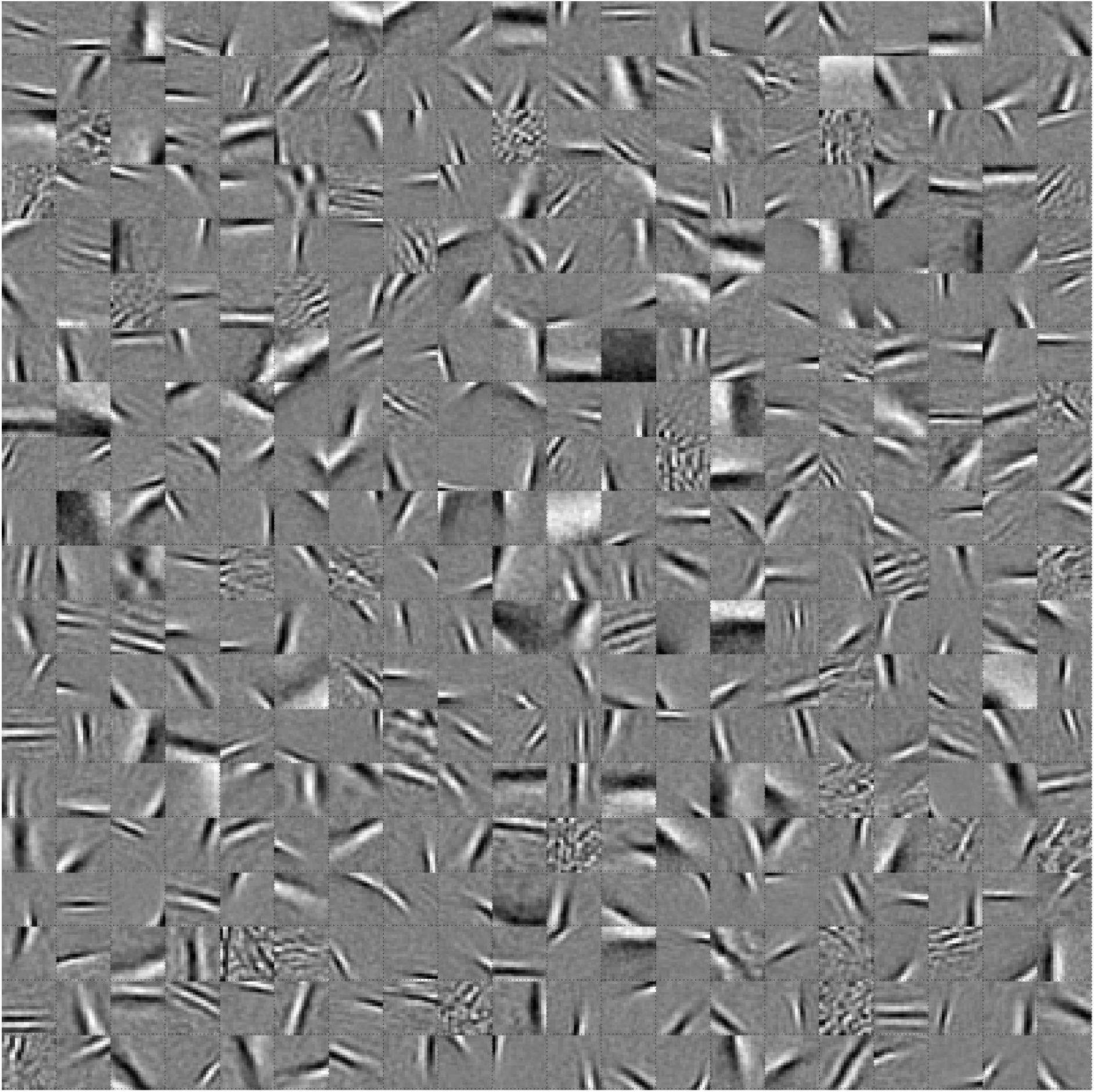
Full set of visual spatial RFs of sparse coding model units trained on visual inputs without added noise. Model units were obtained by training the sparse coding model with 400 hidden units on identical visual inputs used to train the temporal prediction model. The model configuration (400 hidden units, *L*_1_ sparsity strength of 10^0.5^ on the unit activities) was chosen because it resulted in the RFs that look most like the RFs of V1 simple cells as determined by visual inspection. Each square represents the spatial RF of a single unit, shown at its best timestep. The best timestep was determined by selecting the timestep for which the power (sum of squares) of the RF was greatest. White – excitation, black - inhibition.

**Figure 5-Figure Supplement 5.**
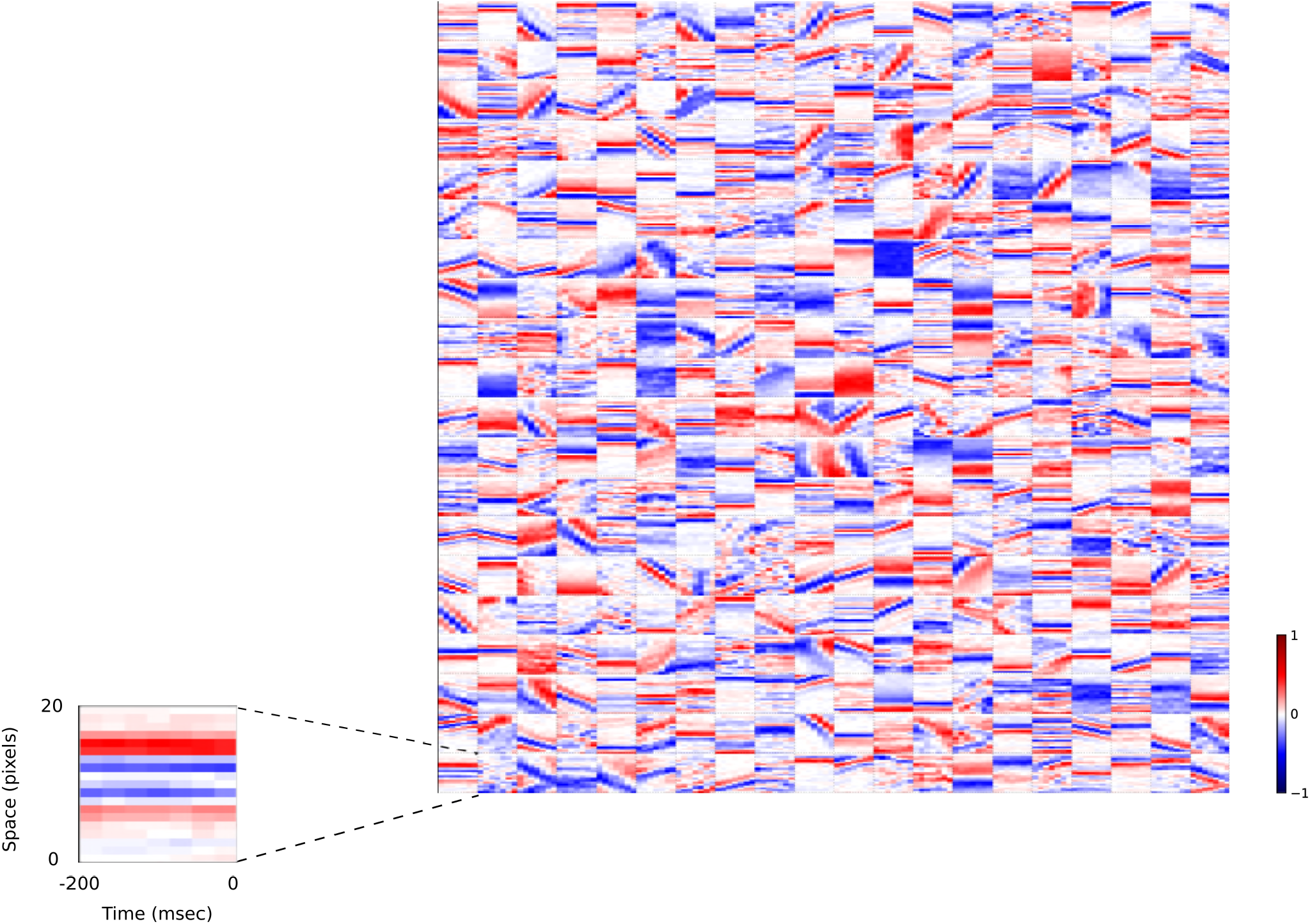
2D (space-time) visual spatiotemporal RFs of sparse coding model units trained on visual inputs without added noise. Obtained from the same units shown in Fig. 5-Fig. Supplement 4 using methods outlined in Fig 2c. Red – excitation, blue - inhibition. Inset shows axes.

**Figure 6-Figure Supplement 1.**
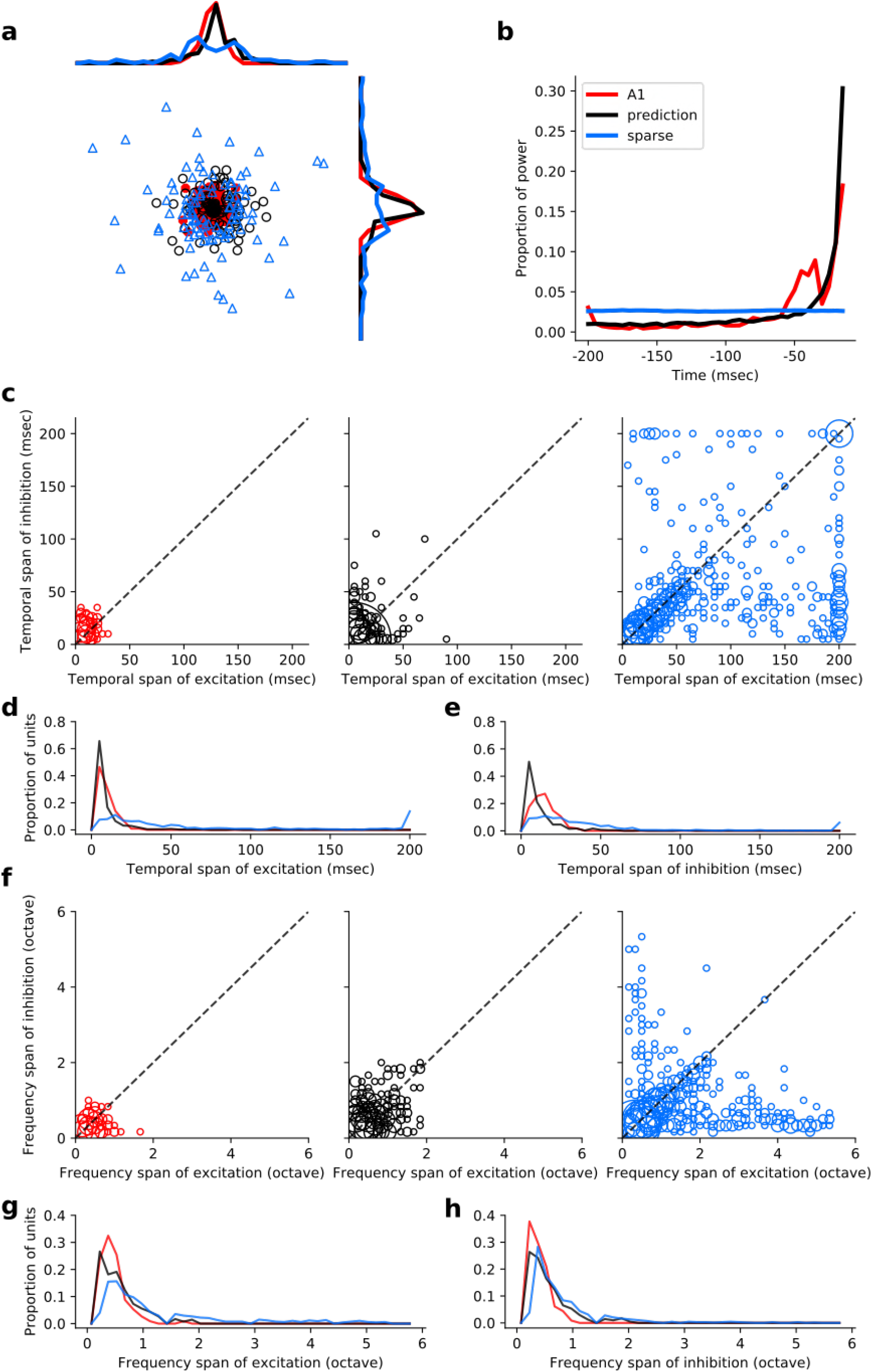
Population measures for real A1 spectrotemporal RFs and temporal prediction and sparse coding model auditory RFs when models are trained on auditory inputs without added noise. Real units are the same as those shown in Figure 3. Temporal prediction model units are the same as those shown in Fig. 4-Fig. Supplement 5; Sparse coding model units are the same as those shown in Fig. 5-Fig. Supplement 1. **a,** Each point represents a single RF (with 32 frequency and 38 time bins) which has been embedded in a 2 dimensional space using Multi-Dimensional Scaling (MDS). Red circles - real A1 neurons, black circles – temporal prediction model units, blue triangles – sparse coding model units. Colour scheme applies to all subsequent panels in Figure. b, Proportion of power contained in each time step of the RF, taken as an average across the population of units. **c,** Temporal span of excitatory subfields versus that of inhibitory subfields, for real neurons and temporal prediction and sparse coding model units. The area of each circle is proportional to the number of occurrences at that point. **d,** Distribution of temporal spans of excitatory subfields, taken by summing along the x-axis in **c. e,** Distribution of temporal spans of inhibitory subfields, taken by summing along the y-axis in **c. f,** Frequency span of excitatory subfields versus that of inhibitory subfields, for real neurons and temporal prediction and sparse coding model units. **g,** Distribution of frequency spans of excitatory subfields, taken by summing along the x-axis in **f. h,** Distribution of frequency spans of inhibitory subfields, taken by summing along the y-axis in **f.**

**Figure 7-Figure Supplement 1.**
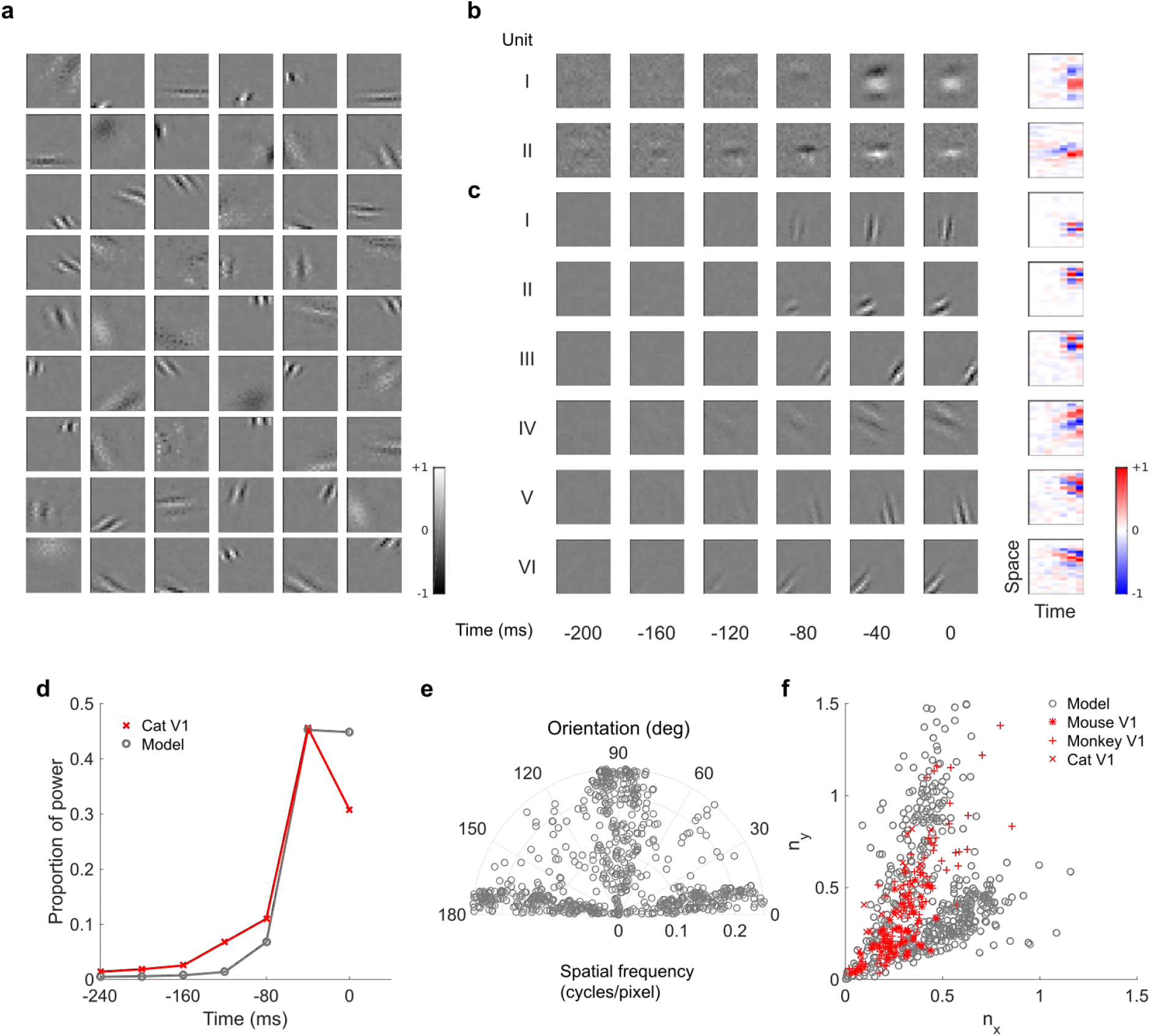
Visual RFs and population measures for real V1 neurons and sparse coding model units. **a,** Model units are the same as those used in Fig. 5-Fig. Supplement 1. Example spatial RFs of randomly selected units at their best timestep. **b-c,** Example 3D and corresponding 2D spatiotemporal RFs at most recent 6 time steps of (**b**) real^23^ V1 neurons and (**c**) sparse coding model units. **d,** Proportion of power (sum of squared weights over space and averaged across units) in each time step, for real and model populations. **e,** Joint distribution of spatial frequency and orientation tuning for population of model units. **f,** Distribution of RF shapes for real neurons (cat^14^, mouse^29^ and monkey^16^) and model units. For **e-f,** only units that could be well approximated by Gabor functions (n = 289 units; see Methods) were included in the analysis. Of these, only model units that were space-time separable (n = 153) are shown in **f** to be comparable with the neuronal data^16^.

**Figure 7-Figure Supplement 2.**
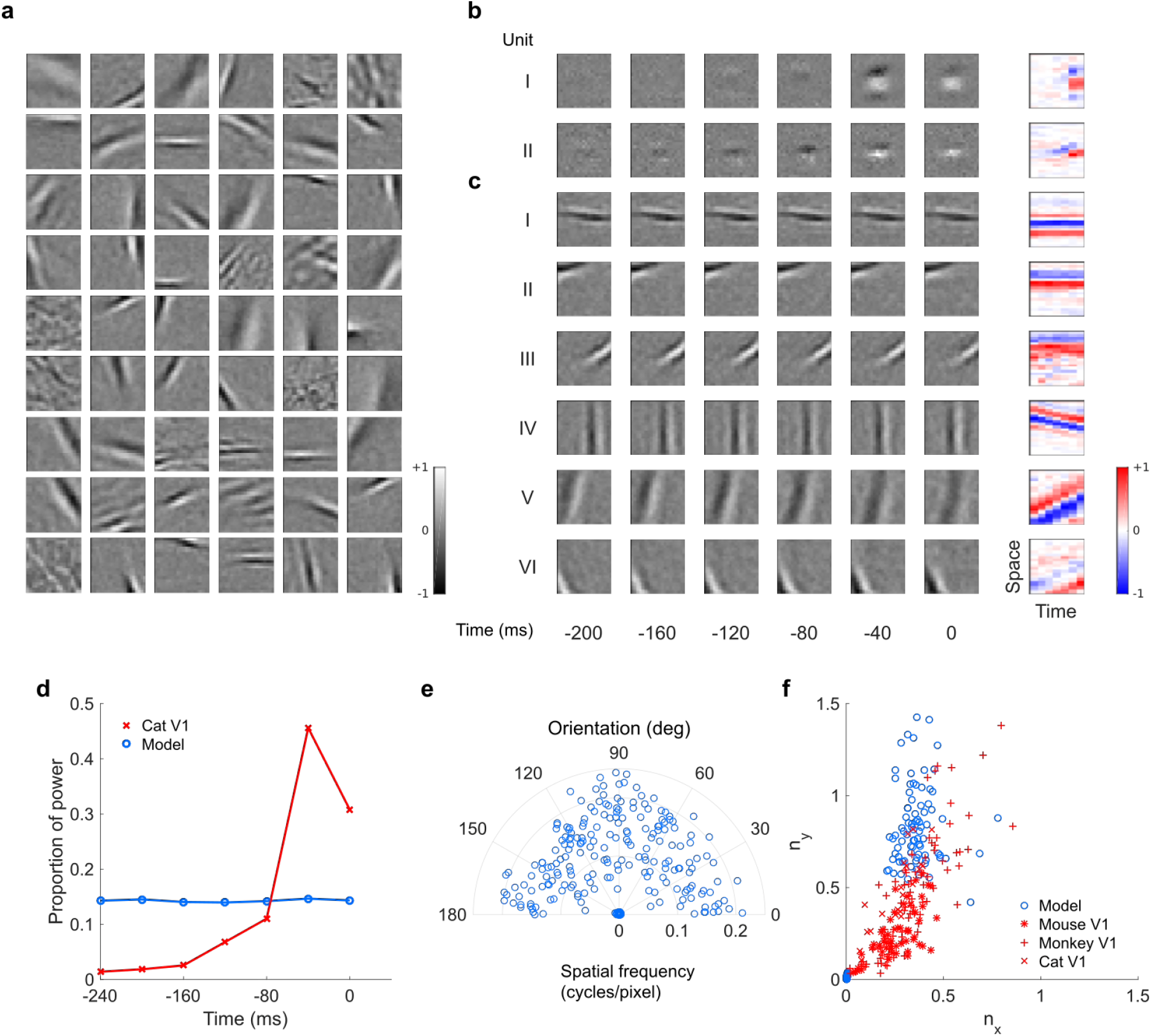
Visual RFs and population measures for real V1 neurons and temporal prediction model units trained on visual inputs without added noise. **a,** Model units are the same as those used in Fig. 11. Example spatial RFs of randomly selected units at their best timestep. **b-c,** Example 3D and corresponding 2D spatiotemporal RFs at most recent 6 time steps of (**b**) real^7^ V1 neurons and (**c**) sparse coding model units. **d,** Proportion of power (sum of squared weights over space and averaged across units) in each time step, for real and model populations. **e,** Joint distribution of spatial frequency and orientation tuning for population of model units. **f,** Distribution of RF shapes for real neurons (cat^4^, mouse^5^ and monkey^6^) and model units. For **e-f,** only units that could be well approximated by Gabor functions (n = 1038 units; see Methods) were included in the analysis. Of these, only model units that were space-time separable (n = 472) are shown in **f** to be comparable with the neuronal data^16^.

**Figure 7-Figure Supplement 3.**
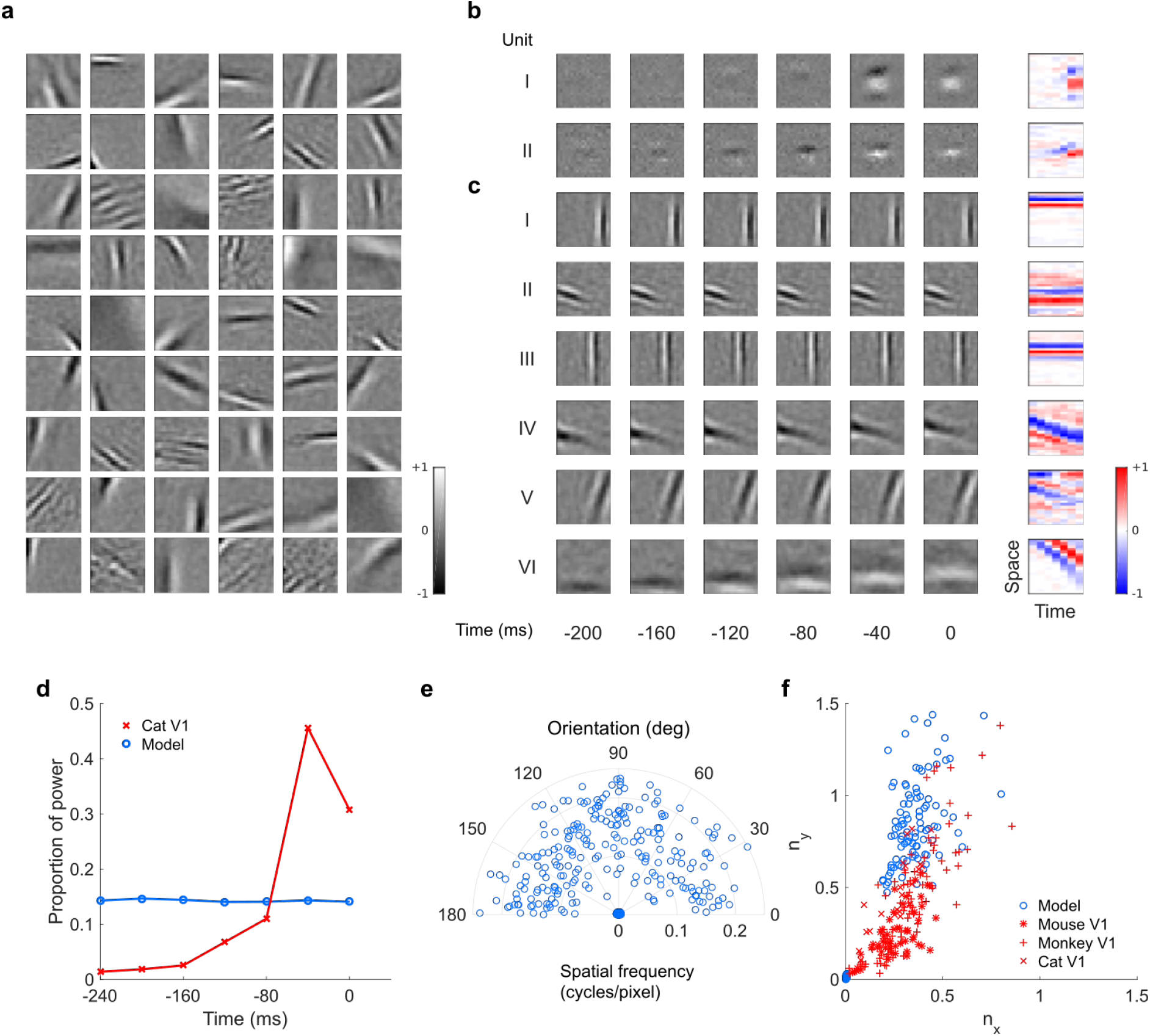
Visual RFs and population measures for real V1 neurons and sparse coding model units trained on visual inputs without added noise. **a,** Model units are the same as those used in Fig. 15. Example spatial RFs of randomly selected units at their best timestep. **b-c,** Example 3D and corresponding 2D spatiotemporal RFs at most recent 6 time steps of (**b**) real^23^ V1 neurons and (**c**) sparse coding model units. **d,** Proportion of power (sum of squared weights over space and averaged across units) in each time step, for real and model populations. **e,** Joint distribution of spatial frequency and orientation tuning for population of model units. **f,** Distribution of RF shapes for real neurons (cat^14^, mouse^29^ and monkey^16^) and model units. For **e-f**, only units that could be well approximated by Gabor functions (n = 323 units; see Methods) were included in the analysis. Of these, only model units that were space-time separable (n = 173) are shown in **f** to be comparable with the neuronal data^16^.

**Figure 8-Figure Supplement 1.**
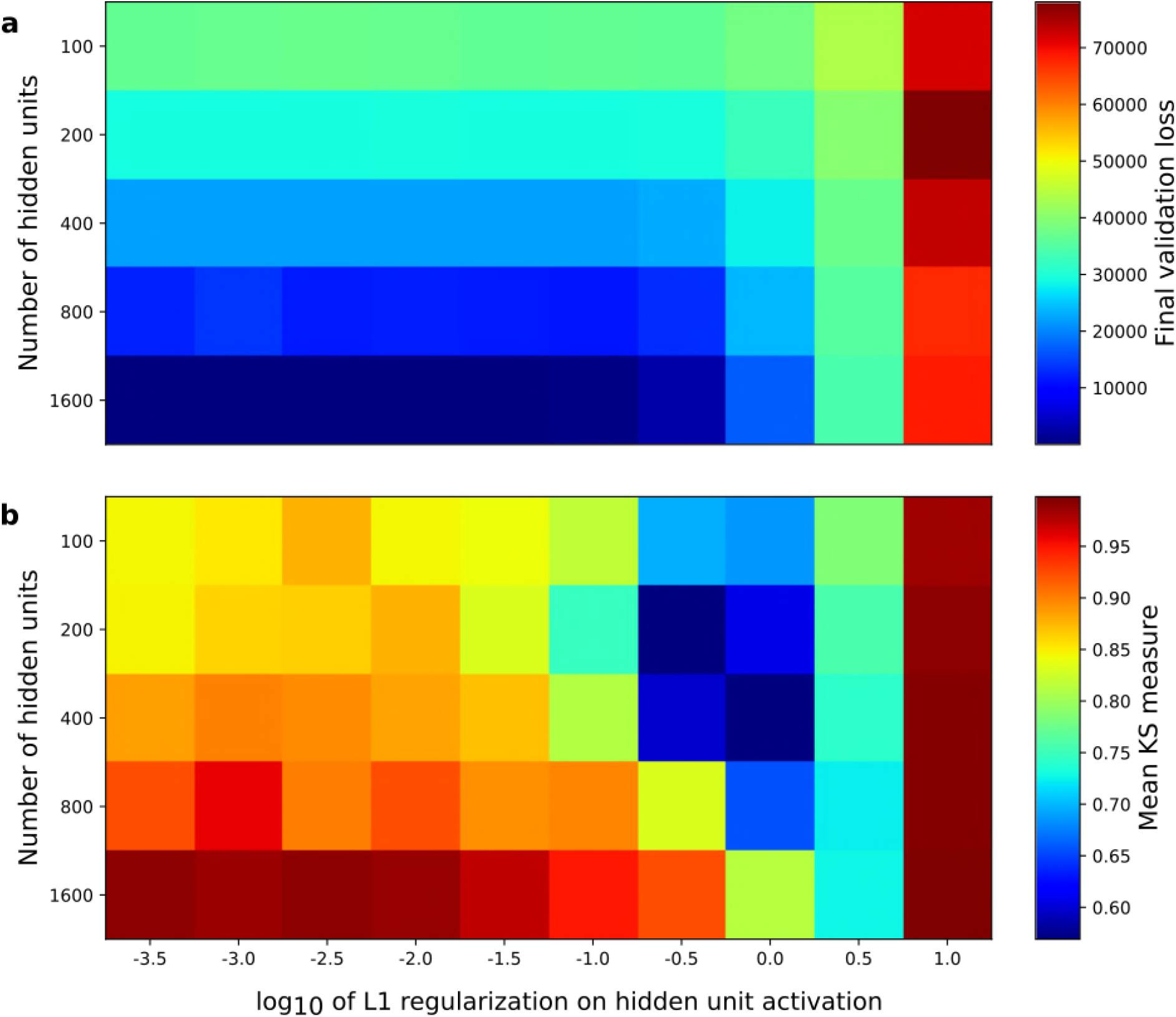
Correspondence between sparse coding model’s ability to reproduce its input and the similarity of its units’ responses to those of real A1 neurons. Performance of model as a function of number of hidden units and regularization on the weights as measured by **a,** reconstruction error on validation set at the end of training and **b,** similarity between model units and real A1 neurons. The similarity between the real and model units is measured by averaging Kolmogorov-Smirnov distance between each of the real and model distributions for the span of temporal and frequency tuning of the excitatory and inhibitory RF subfields (e.g. the distributions in Fig. 3d-e and Fig. 3g-h).

## References

1 Bialek, W., Nemenman, I. & Tishby, N. Predictability, complexity, and learning. Neural Comput. 13, 2409–63 (2001).

2 Attneave, F. Some informational aspects of visual perception. Psychol. Rev. 61, 183–93 (1954).

3 Barlow, H. B. in The Mechanisation of Thought Processes (eds. Blake, D. V. & Uttley, A. M.) 535–539 (H. M. Stationary Office, 1959).

4 Olshausen, B. A. & Field, D. J. Emergence of simple-cell receptive field properties by learning a sparse code for natural images. Nature 381, 607–609 (1996).

5 Olshausen, B. A. & Field, D. J. Sparse coding with an overcomplete basis set: a strategy employed by V1? Vision Research 37, 3311–3325 (1997).

6 van Hateren, J. H. & Ruderman, D. L. Independent component analysis of natural image sequences yields spatio-temporal filters similar to simple cells in primary visual cortex. Proc.R. Soc. Lond. B 265, 2315–2320 (1998).

7 Berkes, P. & Wiskott, L. Slow feature analysis yields a rich repertoire of complex cell properties. J. Vis. 5, 579–602 (2005).

8 Berkes, P., Turner, R. E. & Sahani, M. A structured model of video reproduces primary visual cortical organisation. PLoS Comput. Biol. 5, (2009).

9 Klein, D. J., König, P. & Körding, K. P. Sparse spectrotemporal coding of sounds. EURASIP J. Appl. Signal Processing 2003, 1–9 (2003).

10 Carlson, N. L., Ming, V. L. & DeWeese, M. R. Sparse codes for speech predict spectrotemporal receptive fields in the inferior colliculus. PLoS Comput. Biol. 8, (2012).

11 Zhao, L. & Zhaoping, L. Understanding auditory spectro-temporal receptive fields and their changes with input statistics by efficient coding principles. PLoS Comput. Biol. 7, (2011).

12 Kozlov, A. S. & Gentner, T. Q. Central auditory neurons have composite receptive fields. Proc. Natl. Acad. Sci. 113, 1441–1446 (2016).

13 Cusack, R. & Carlyon, R. in Echological pyschoacoustics (ed. Neuhoff, J. G.) 15–48 (Elsevier, 2004).

14 Hubel, D. H. & Wiesel, T. N. Receptive fields of single neurones in the cat’s striate cortex. J. Physiol. 148, 574–91 (1959).

15 Jones, J. P. & Palmer, L. A. An evaluation of the two-dimensional Gabor filter model of simple receptive fields in cat striate cortex. J. Neurophysiol. 58, 1233–1258 (1987).

16 DeAngelis, G. C., Ohzawa, I. & Freeman, R. D. Spatiotemporal organization of simple-cell receptive fields in the cat’s striate cortex. I. General characteristics and postnatal development.J. Neurophysiol. 69, 1091–1117 (1993).

17 Ringach, D. L. Spatial structure and symmetry of simple-cell receptive fields in macaque primary visual cortex. J. Neurophysiol. 88, 455–463 (2002).

18 deCharms, R. C., Blake, D. T. & Merzenich, M. M. Optimizing sound features for cortical neurons. Science 280, 1439–43 (1998).

19 van Hateren, J. H. & van der Schaaf, A. Independent component filters of natural images compared with simple cells in primary visual cortex. Proc. R. Soc. London B 265, 359–366 (1998).

20 Eliasmith, C. & Anderson, C. H. Neural engineering: computation, representation, and dynamics in neurobiological systems. (MIT Press, 2003).

21 Willmore, B. D. B., Schoppe, O., King, A. J., Schnupp, J. W. H. & Harper, N. S. Incorporating midbrain adaptation to mean sound level improves models of auditory cortical processing. J. Neurosci. 36, 280–9 (2016).

22 Carlin, M. A. & Elhilali, M. Sustained firing of model central auditory neurons yields a discriminative spectro-temporal representation for natural sounds. PLoS Comput. Biol. 9, (2013).

23 Brito, C. S. N. & Gerstner, W. Nonlinear Hebbian learning as a unifying principle in receptive field formation. PLoS Comput. Biol. 12, (2016).

24 Ohzawa, I., DeAngelis, G. C. & Freeman, R. D. Encoding of binocular disparity by simple cells in the cat’s visual cortex. J. Neurophysiol. 75, 1779–805 (1996).

25 Olshausen, B. A. Learning sparse, overcomplete representations of time-varying natural images. IEEE Int. Conf. Image Process. (2003).

26 Hyvärinen, A., Hurri, J. & Väyrynen, J. Bubbles: a unifying framework for low-level statistical properties of natural image sequences. J. Opt. Soc. Am. A 20, 1237–1252 (2003).

27 Młynarski, W. & McDermott, J. H. Learning Mid-Level Auditory Codes from Natural Sound Statistics. (2017). at <http://arxiv.org/abs/1701.07138>

28 Blättler, F., Hahnloser, R. H. R., Doupe, A., Hahnloser, R. & Wilson, R. An Efficient Coding Hypothesis Links Sparsity and Selectivity of Neural Responses. PLoS One 6, e25506 (2011).

29 Kreile, A. K., Bonhoeffer, T. & Hübener, M. Altered visual experience induces instructive changes of orientation preference in mouse visual cortex. J. Neurosci. 31, 13911–13920 (2011).

30 Niell, C. M. & Stryker, M. P. Highly Selective Receptive Fields in Mouse Visual Cortex. J. Neurosci. 28, (2008).

31 Attneave, F. Some informational aspects of visual perception. Psychol. Rev. 61, 183–93 (1954).

32 Srinivasan, M. V, Laughlin, S. B. & Dubs, A. Predictive coding: a fresh view of inhibition in the retina. Proc. R. Soc. London. Ser. B, Biol. Sci. 216, 427–59 (1982).

33 Palmer, S. E., Marre, O., Berry, M. J. & Bialek, W. Predictive information in a sensory population. Proc. Natl. Acad. Sci. U. S. A. 112, 6908–13 (2015).

34 Salisbury, J. M. & Palmer, S. E. Optimal prediction in the retina and natural motion statistics.J. Stat. Phys. 162, 1309–1323 (2016).

35 Huang, Y. & Rao, R. P. N. Predictive coding. Wiley Interdiscip. Rev. Cogn. Sci. 2, 580–593 (2011).

36 Smith, E. C. & Lewicki, M. S. Efficient auditory coding. Nature 439, 978–982 (2006).

37 Wiskott, L. & Sejnowski, T. J. Slow feature analysis: unsupervised learning of invariances. Neural Comput. 14, 715–770 (2002).

38 Creutzig, F. & Sprekeler, H. Predictive coding and the slowness principle: an information-theoretic approach. Neural Comput. 20, 1026–1041 (2008).

39 Palm, R. B. Prediction as a candidate for learning deep hierarchical models of data. (Technical University of Denmark, (DTU) Informatics, 2012).

40 Rao, R. P. & Ballard, D. H. Predictive coding in the visual cortex: a functional interpretation of some extra-classical receptive-field effects. Nat. Neurosci. 2, 79–87 (1999).

41 Friston, K. Learning and inference in the brain. Neural Networks 16, 1325–1352 (2003).

42 Marzen, S. E. & DeDeo, S. The evolution of lossy compression. J. R. Soc. Interface 14, (2017).

43 Field, D. J. Relations between the statistics of natural images and the response properties of cortical cells. J. Opt. Soc. Am. A 4, 2379 (1987).

44 Torralba, A. & Oliva, A. Statistics of natural image categories. Comput. Neural Syst 14, 391–412 (2003).

45 Rees, A. & Møller, A. R. Responses of neurons in the inferior colliculus of the rat to AM and FM tones. Hear. Res. 10, 301–330 (1983).

46 Harris, K. D. & Mrsic-Flogel, T. D. Cortical connectivity and sensory coding. Nature 503, 51–58 (2013).

47 Srivastava, N., Mansimov, E. & Salakhutdinov, R. Unsupervised learning of video representations using LSTMs. (2015). at <http://arxiv.org/abs/1502.04681>

48 Ranzato, M. et al. Video (language) modeling: a baseline for generative models of natural videos. (2016). at <http://arxiv.org/abs/1412.6604>

49 Lotter, W., Kreiman, G. & Cox, D. Deep Predictive Coding Networks for Video Prediction and Unsupervised Learning. (2016). at <http://arxiv.org/abs/1605.08104>

50 Oh, J., Guo, X., Lee, H., Lewis, R. & Singh, S. Action-Conditional Video Prediction using Deep Networks in Atari Games. Adv. Neural Inf. Process. Syst. 28, (2015).

51 Rubin, J., Ulanovsky, N., Nelken, I. & Tishby, N. The representation of prediction error in auditory cortex. PLoS Comput. Biol. 10, 1–28 (2016).

52 Morrison, G. S. et al. Forensic database of voice recordings of 500+ Australian English speakers. (2015). at <http://databases.forensic-voice-comparison.net/>

53 Morrison, G. S., Rose, P. & Zhang, C. Protocol for the collection of databases of recordings for forensic-voice-comparison research and practice. Aust. J. Forensic Sci. 44, 155–167 (2012).

54 Sachs, M. B. & Abbas, P. J. Rate versus level functions for auditory-nerve fibers in cats: tone-burst stimuli. J. Acoust. Soc. Am. 56, 1835–47 (1974).

55 Kingma, D. P. & Ba, J. Adam: A Method for Stochastic Optimization. (2014). at <http://arxiv.org/abs/1412.6980>

56 Sohl-Dickstein, J., Poole, B. & Ganguli, S. Fast large-scale optimization by unifying stochastic gradient and quasi-Newton methods. Proc. 31st Int. Conf. Mach. Learn. 32, 604–612 (2014).

57 Simoncelli, E., Pillow, J. W., Paninski, L. & Schwartz, O. in The cognitive neurosciences, III (ed. Gazzaniga, M.) 327–338 (MIT Press, 2004).

58 Dahmen, J. C., Hartley, D. E. H. & King, A. J. Stimulus-Timing-Dependent Plasticity of Cortical Frequency Representation. 28, 13629–13639 (2008).

59 Craig A. Atencio, Tatyana O. Sharpee, and C. E. S. Nonlinearities in Auditory Cortical Neurons. 58, 956–966 (2009).

60 Chichilnisky, E. J. A simple white noise analysis of neuronal light responses. Network 12, 199–213 (2001).

61 Rabinowitz, N. C., Willmore, B. D. B., Schnupp, J. W. H. & King, A. J. Contrast Gain Control in Auditory Cortex. Neuron 70, 1178–1191 (2011).

62 Sahani, M. & Linden, J. F. J. F. How linear are auditory cortical responses? Adv. Neural Inf. Process. Syst. 15, 109–116 (2003).

